# Cilia beating of ependymal cells regulates adult neural stem cell quiescence via mechanical forces mediated by PKD1/2-TRPM3

**DOI:** 10.64898/2026.02.26.708348

**Authors:** Cedric Bressan, Archana Gengatharan, Raquel Rodriguez-Aller, Maria L. Richter, Marina Snapyan, Judith Fischer-Sternjak, Maryam Rezaeezadeh Roukerd, Nicole Rosin, Abigael Cherinet, Jeff Biernaskie, Ehsan Habibi, Magdalena Götz, Armen Saghatelyan

## Abstract

In many tissues stem cells are located lining a fluid-filled volume and their neighboring niche cells include cells with beating cilia. However, the role of mechanical forces created by cilia beating on stem cells remains elusive. We developed an approach to transiently inhibit the cilia beating of ependymal cells (EC) lining the forebrain ventricle by injecting magnetic beads-coupled antibodies targeting EC cilia and then applying a magnetic field. We show that EC cilia beating enforces neural stem cells (NSCs) quiescence through mechano-sensitive PKD1/2- and TRPM3-mediated Ca^2+^ transients. Only a few hours of EC cilia beating inhibition triggered NSC activation *in vivo*. CRISPR-Cas9-mediated deletion of TRPM3 or PKD1/2 in NSCs phenocopied the effect of EC cilia beating inhibition, while TRPM3 pharmacological activation rescued NSC quiescence in the absence of cilia beating. Our data reveal a novel regulator of stem cells exposed to fluids via the mechanical forces mediated by cilia beating.

## Introduction

Somatic stem cells located in various organs are often exposed to a lumen filled with fluid and are surrounded by cells with multiple beating cilia. In the lungs, gut, kidneys and specific regions of embryonic and adult brains these multi-ciliated cells play vital functions by regulating fluid flow and by circulating various factors that affect somatic stem cells functions through diverse signaling mechanisms^1-9^. Alterations in the formation and/or function of cilia lead to ciliopathy and result in major developmental disorders^10-12^. Cilia beating, however, not only circulates factors and regulates fluids flow, but may also exert a mechano-sensitive influence on the physiology of distinct somatic stem cells. Although mechanical forces are increasingly becoming recognized as major regulators of cell structure and function^13,14^, little is known about their effects on stem cells. In addition, it is not known whether cilia beating directly affects stem cell functions.

In the adult brain, neural stem cells (NSCs) residing in the subventricular zone (SVZ) are exposed to a variety of factors from the cerebral spinal fluid (CSF) via small apical processes bearing a non-motile primary cilium^2,7,15,16^. NSCs are surrounded by multi-ciliated ependymal cells (ECs) that are organized in a pinwheel structure around the apical process of the NSCs^16-19^. Several studies have shown that factors in the CSF affect adult NSC function^2,7,20,21^. It has been also shown *in vitro* that modulating the mechano-sensitive properties of NSCs themselves, either by manipulating the stiffness of the substrate or by applying sheer stress, affects their proliferation and differentiation^22-24^. These results are in line with those of loss-of-function experiments for specific mechano-sensitive receptors, ion channels, and downstream signaling cascades in NSCs that affect their proliferation and differentiation^22-24^. These studies provided important insights into the roles of some mechano-sensitive channels/receptors in NSCs and how mechanical forces applied either directly on NSCs *in vitro* or on entire brain sections affect NSC proliferation and differentiation. However, the endogenous source of these mechanical forces *in vivo* remains unclear. Likewise, the extent and effects of the mechanical forces exerted by EC cilia beating on NSC sensory cilia are unknown.

To address these questions, we developed an approach to acutely and selectively modulate EC cilia beating, while monitoring the functional activity of adult NSCs and their quiescence/activation dynamics. We inhibited cilia beating by targeting EC cilia with an EC-specific cell surface antibody directed against CD24 coupled to magnetic beads. We then applied a magnetic field, both *ex vivo* and *in vivo,* to acutely and transiently inhibit cilia beating. This revealed that the mechanical forces created by EC cilia beating were required for the maintenance of the quiescent state of NSCs and that halting cilia beating triggered NSC activation linked to increased protein translation. The EC cilia beating-mediated effect on NSCs was dependent on Ca^2+^ permeable TRPM3 receptors being enriched in NSCs as well as mechano-sensitive polycystin 1/2 (PKD1/2) channels. EC cilia beating inhibition or CRISPR-Cas9-mediated deletion of TRPM3 or PKD1/2 from NSCs abrogated the Ca^2+^ dynamics required for the maintenance of the quiescent state and resulted in NSC activation, as observed after the inhibition of ECs cilia beating. Conversely, pharmacological activation of TRPM3 fully reverted the effects of reduced cilia beating on NSCs *in vivo*. Altogether, our data indicate that EC cilia beating has crucial effects on the state of adult NSCs through mechano-sensitive receptors and downstream Ca^2+^ signaling, and that the abrogation of cilia beating triggers NSC activation. These data thus point to a novel regulatory mechanism of NSC activation with key physiological relevance, as cilia beating is affected in several physiological and pathological conditions^10,12,25-29^.

## Results

### Acute electro-magnetic arrest of ECs cilia beating

To influence adult EC cilia beating, both *ex vivo* in acute adult brain sections and *in vivo*, we developed an approach that made it possible to selectively couple magnetic beads to EC cilia and to apply a magnetic field to reduce their beating (**Fig. 1**). To do so, we coupled antibodies directed against the CD24 antigen, which is expressed on EC cilia, to magnetic beads (**Fig. 1A**). As a control, we used magnetic beads coupled to antibodies against PSA-NCAM or PDGFRα, which are not expressed by EC. As CD24 may also be expressed in immature neurons and glia^30,31^, the antibodies were injected into the lateral ventricle (LV) of adult mice, leading to the selective binding of CD24-coupled, but not PSA-NCAM-coupled or PDGFRα-coupled, antibodies to the cilia of ECs thus avoiding the binding of CD24-coupled antibodies to immature neurons and glia (**Fig. 1B and Supp. Fig. 1A**). The beads were visualized using either visible light (**Fig. 1B**) or 564 nm excitation (**Fig. 1C**), either in sagittal (**Fig. 1B**) or coronal (**Fig. 1C**) sections, the latter to further ascertain the presence of beads specifically in the LV. To track the beating of beads coupled to EC cilia, we prepared acute brain sagittal sections 30-45 min after the injection of the beads into the LV and performed time-lapse imaging of bead movement with an acquisition rate of 80-100 frames/s (**Fig. 1D**). Our experiments revealed reliable and fast movement of either individual or clustered beads, the latter likely reflecting the binding of multiple beads to the cilia of the same multiciliated EC (**Fig. 1D**). To quantify the frequency of bead movement, we tracked the centroid of the bead in all consecutive frames, which gave a beating frequency of 23.2 ± 0.5 Hz (n=47 beads from 10 mice; **Fig. 1E-H**). To ascertain that beads binding to the cilia did not affect their beating frequency, we assessed the beating frequency of fluorescently labeled EC cilia. To do so, we electroporated a plasmid encoding a membrane-tethered GFP (CMV-Lck-GFP) that was injected into the LV of adult 1-2-month-old mice, resulting in labeling of ECs (**Fig. 1E-F**). Time-lapse imaging of cilia in acute brain slices 3-5 days after electroporation revealed that the frequency of beating of fluorescently labeled ECs cilia (24.3 ± 0.7 Hz) was the same as that of those coupled to magnetic beads (23.2 ± 0.5 Hz; **Fig. 1F-H**), indicating that the binding of the beads did not affect the beating frequency of the cilia. We next inhibited EC cilia beating by delivering a magnetic field via an electrical current-carrying copper wire positioned in proximity to the surface of an acute brain section covering a small part of the SVZ (**Fig. 1I-J**). The beating frequency of the beads was imaged under baseline conditions and following the delivery of a 2.8-mT electro-magnetic field. Interestingly, our experiments indicated that delivering a 2.8-mT 5-s pulse completely arrested cilia beating (23.1 ± 0.6 Hz under baseline condition vs 0 Hz after magnetic stimulation; n=41 beads from 9 mice; **Fig. 1K-L**). The results of these experiments show that the binding of magnetic beads to EC cilia did not affect their beating frequency, which can be acutely modulated by the delivery of an electro-magnetic field. This opened the possibility of studying the mechano-sensitive role of EC cilia beating on the physiology and quiescence/activation dynamics of adult NSCs that are surrounded by ECs.

**Figure 1:**
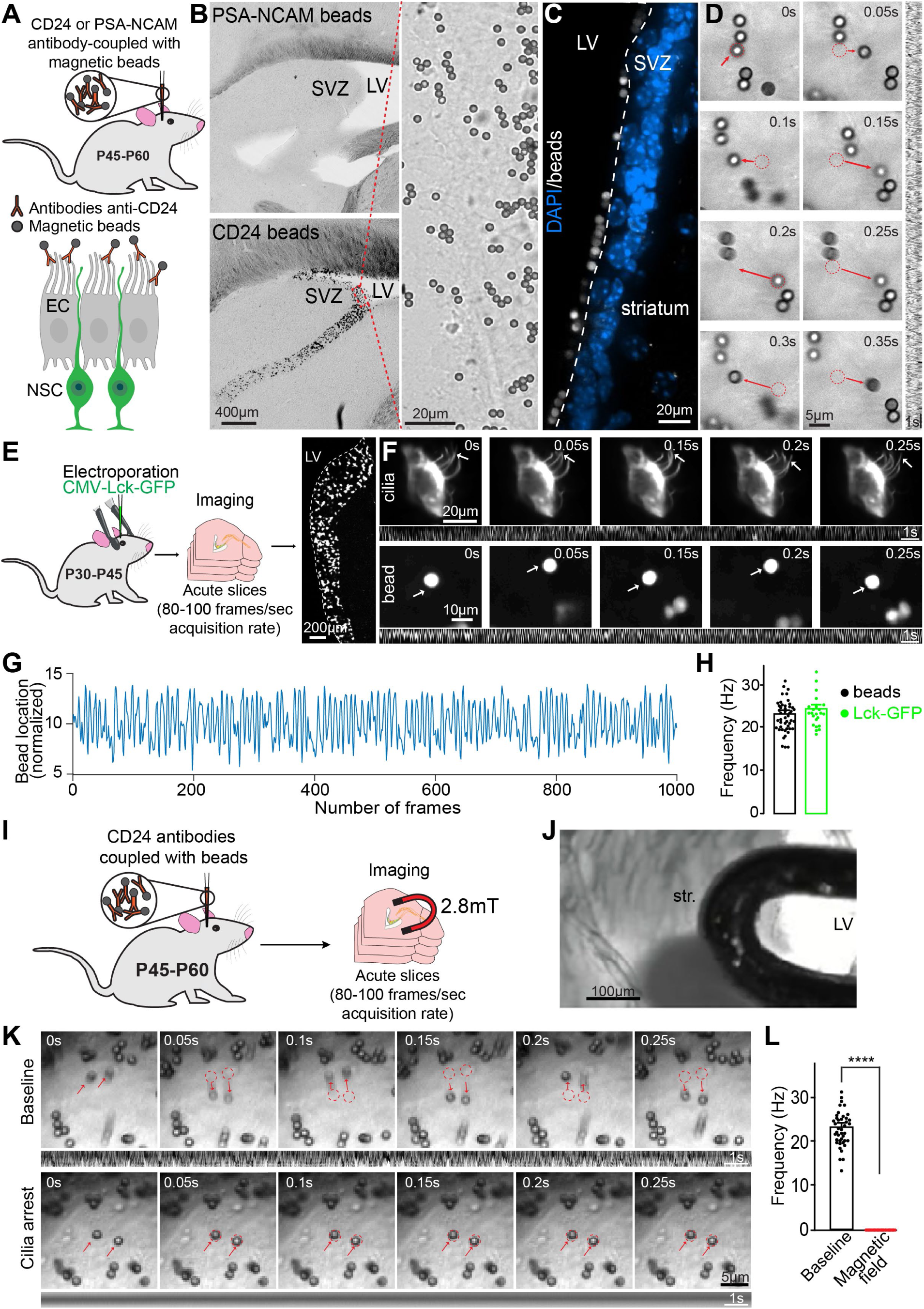
Electro-magnetic modulation of EC cilia beating. **A:** Experimental approach for modulating EC cilia beating consisting of coupling magnetic beads to CD24 antibodies and injecting them into the LV of adult mice. CD24 is expressed by ECs and is localized at their cilia. As a control, we used PSA-NCAM or PDGFRα antibodies. These antigens are not expressed by ECs. **B:** Sagittal sections of adult mouse brains showing the binding of beads coupled to CD24 antibodies to ECs lining the wall of the LV. Each black dot indicates a CD24-coupled bead. Note the lack of beads coupled to the PSA-NCAM antibodies following their injection into the LV. The right image shows a high magnification of CD24 antibody-coupled beads. **C:** Coronal section of an adult mouse brain showing the specific binding of magnetic beads coupled to CD24 antibodies to EC cilia and the presence of beads in the LV. The beads were imaged at 564nm excitation wavelength, and the slices were counterstained with DAPI. **D:** Time-lapse imaging and kymograph of beads movement in acute brain sections prepared from mice in which CD24 antibody-coupled beads were injected into the LV. The imaging was performed 1-4 h after the injection of the beads into the LV with an acquisition rate of 80-100 frames/s. The red ROI indicate the positions of the beads at the previous time-point. **E:** Schematic drawing of the electroporation approach used to label ECs with membrane-tethered GFP (Lck-GFP), the time-lapse imaging of acute brain slices, and image showing electroporated cells in the adult SVZ. **F:** Time-lapse imaging of EC cilia labeled either with membrane-tethered GFP (upper panels) or magnetic beads (lower panels). The bottom panels show the kymographs. **G:** An example of bead tracking based on the assessment of the centroid in each consecutive time-point during 12.5 s of imaging. **H:** Quantification of the frequency of cilia beating assessed by time-lapse imaging of cilia labeled either with Lck-GFP or magnetic beads. **I:** Experimental approach for arresting EC cilia beating *ex vivo* in acute brain slices. Beads coupled to CD24 antibodies were injected into the LV of adult mice, and acute brain slices were prepared 30-60 min after injection. The electro-magnetic field was applied through a coiled copper wire positioned at the surface of the slice. **J:** Photomicrograph showing an acute brain slice and a wire used for the application of the electro-magnetic field. **K-L:** Time-lapse imaging, kymograph (bottom panels) and quantification of bead movement in acute brain sections under baseline conditions (upper row) and after the application of a 2.8-mT magnetic field (lower row). The red ROIs indicate the position of the beads at the previous time-point. Statistical significance was determined using a paired Student’s t-test. \*\*\*\**p*<0.0001. Data are expressed as means ± SEM. N=24 Lck-GFP+ cilia and 48 beads from 3 and 10 mice, respectively, for **H**, and 41 beads from 9 mice for **L**. See also **Supplementary Figure 1**.

### *In vivo* inhibition of EC cilia beating triggers NSC activation

To modulate cilia beating *in vivo*, we injected CD24- or PSA-NCAM antibodies coupled with magnetic beads into the LV of adult mice that had previously been electroporated with GFAP-GFP and prominin (Prom)-mCherry plasmids at P0-P1. As the coincident activity of the GFAP and Prom promoters is present only in NSCs and not in other cell types in the SVZ niche^32-35^, this approach made it possible to unequivocally label NSCs and track their activation. Thirty to 45 min after the beads were injected into the LV and the animals had recovered after the stereotaxic injections, the mice were placed in a custom-built behavioral chamber consisting of two cages connected with a tunnel (**Fig. 2A-B**). Permanent neodymium magnets were placed on each side of the tunnel, which resulted in about 300-mT magnetic field inside the tunnel (**Fig. 2A-B)**. No magnetic field was detected in the cages (**Fig. 2A-B**). Each time a mouse passed through the tunnel from one cage to another it was exposed to the magnetic field (**Fig. 2B**). The use of a narrow tunnel and 4-inch-long magnets positioned on the sides of the tunnel made it possible to ensure that the mice injected with either with CD24- or PSA-NCAM antibodies coupled with magnetic beads were exposed to a magnetic field of similar strength, direction and angle. In contrast to the administration of a magnetic field administration in the home-cage where the strength and direction of the magnetic field perceived by the mouse might vary depending on the position of the mouse in the cage and its behavior (rearing, running, resting, etc), the delivery of the magnetic field in a narrow tunnel where mice could only move from one cage to another ensured the consistency of the strength, direction and angle of the magnetic field applied to each and every mouse. We left the mice in these behavioral cages for 3 h and monitored the number of entries and the time that they spent in the tunnel. Immediately after the 3-h exploration period, we prepared acute brain slices and imaged the beating of the beads coupled to EC cilia (**Fig. 2C**). Our analyses revealed that there was a marked decrease in the frequency of cilia-coupled beads, indicating that EC cilia beating *in vivo* was efficiently decreased (22.2 ± 0.5 Hz for mice injected with CD24 antibody-coupled beads but not exposed to the magnets, vs. 5.6 ± 0.8 Hz for mice injected with CD24 antibody-coupled beads and exposed to the magnets; **Fig. 2D**). This effect was transient, given that exposing mice injected with CD24 antibody-coupled beads to the magnetic field for 3 h, followed by a 3 h recovery period during which the mice were housed in their home cage and were not exposed to the permanent magnets, resulted in normal beating of EC cilia coupled to magnetic beads in the acute brain slices (23.9 ± 0.3 Hz for mice injected with CD24 antibody-coupled beads exposed to the permanent magnets for 3 h followed by a 3 h recovery period; **Fig. 2C-D**). Interestingly, the inhibition of EC cilia beating did not affect the overall CSF flow that we assessed by *in vivo* imaging of fluorescent dextran diffusion, injected immediately after inhibiting EC cilia beating (**Supp. Fig. 1**). The mice injected with CD24 antibody-coupled magnetic beads and either exposed or not exposed to the permanent magnet showed the same spread of fluorescent dextran to the contralateral hemisphere (**Supp. Fig. 1**). These data suggested that while EC cilia beating inhibition might affect the local flow of CSF in the bead-occupied areas, the global flow was sustained, likely due to the beating of EC cilia not coupled to magnetic beads.

**Figure 2.**
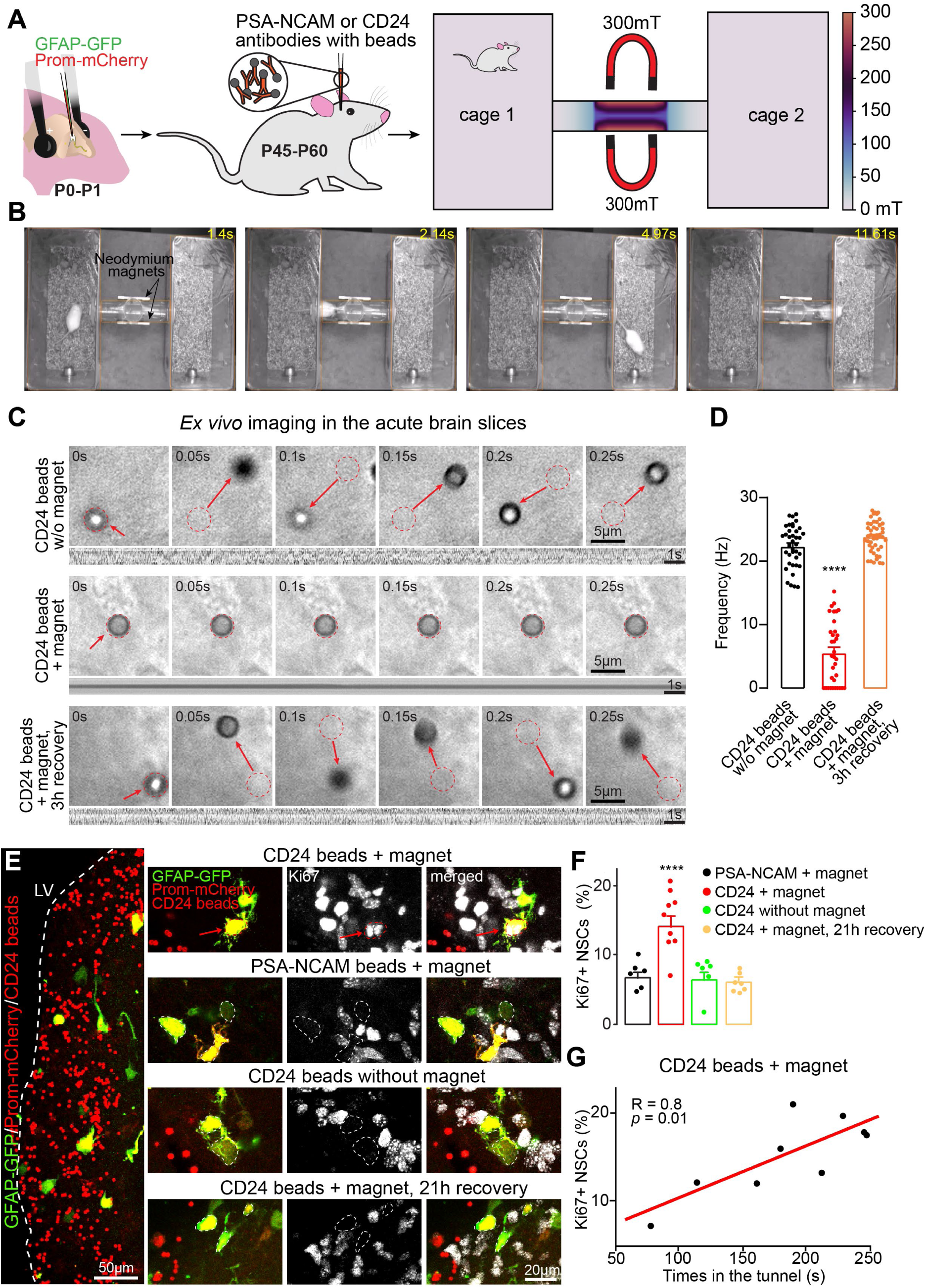
Decrease in EC cilia beating triggers NSC activation. **A-B:** Schematic of the electroporation approach of the GFAP-GFP and Prom-mCherry plasmids used to label NSCs, and CD24 antibody-coupled (or as a control PSA-NCAM antibody-coupled) bead injections into the LV to modulate EC cilia beating *in vivo*. Thirty to 45 min after the injection of the beads, the mice were placed in a behavioral chamber consisting of two cages connected by a tunnel equipped with permanent neodymium magnets. The mice freely explored the behavioral chamber for 3 h, and each time they passed through the tunnel they were exposed to the magnetic field. There was no magnetic field inside either cage. The panel also shows the heatmap of magnetic field measurements in the behavioral cages. **B:** Video imaging of mouse behavior in the two cages connected by a tunnel equipped with permanent neodymium magnets. **C:** Time-lapse imaging and kymographs (bottom panels) of beads in acute brain slices prepared from mice exploring the behavioral chambers shown in **A-B**. The slices were prepared either right after the 3-h period that the animal spent in the behavioral chambers (upper and middle panels) or after an additional 3-h recovery period (3 h magnet +3 h recovery; lower panel). The red ROIs indicate the positions of the beads in each preceding time-point of the time-lapse imaging. **D:** Quantification of the EC cilia beating frequency. Note the marked reduction in the frequency of cilia beating right after the 3-h period that the mice spent in the behavioral chambers (red dots) and the restoration of EC cilia beating when the mice were housed for 3 h in their home cages after being exposed to magnetic field stimulation (orange dots). As a control, we used animals that had been injected with CD24 antibody-coupled beads and that explored the same behavioral chambers for 3 h but without permanent magnets in the tunnel (black dots). Each dot represents individual CD24 antibody-coupled bead that was tracked for 30 s. **E:** Low and high magnification images showing immunolabeling for GFP, mCherry, and Ki67 to assess the percentage of activated NSCs under control conditions and following the arrest of ECs cilia beating. The red arrows and ROI indicate Ki67+ NSCs while the white ROI indicate Ki67-NSCs. Note, the presence of beads in mice injected with CD24 antibody-coupled beads. Only NSCs surrounded by beads coupled to EC cilia were analyzed. **F:** Quantification of the percentage of activated Ki67+ NSCs in different conditions. Note the marked increase in the percentage of activated NSCs following the reduction in the EC cilia beating frequency compared to the two control conditions. Note also that the effect on NSC proliferation was transient as evidenced by the quantification of Ki67+ NSCs in mice exposed to the magnet for 3 h followed by a 21-h recovery period. **G:** Correlation between the time spent in the tunnel and the percentage of activated NSCs following the reduction in the EC cilia beating frequency. Data are expressed as means ± SEM. Statistical significance was determined using a one-way ANOVA followed by a post hoc Fisher’s LSD test for **D** and **F**. \*\*\**p*<0.001. The individual values of the experiments (circles) are also shown. For **D,** n=39, 34, and 48 CD24 antibody-coupled beads from 5, 6, and 4 mice, respectively. For **F**, 640, 876, 516 and 1149 NSCs were analyzed for the PSA-NCAM+magnet (6 mice), CD24+magnet (9 mice), CD24 without magnet (6 mice) and CD24+magnet+21-h recovery (9 mice) conditions, respectively.

We next determined whether the acute and transient reduction in EC cilia beating affected NSC quiescence/activation by quantifying Ki67+ proliferative GFAP-GFP+/Prom-mCherry+ NSCs (**Fig. 2E**). Our analyses revealed that the percentage of activated Ki67+ NSCs in mice injected with CD24 antibody-coupled beads and exposed to the magnetic field increased 2-fold (15.3 ± 1.5%; **Fig. 2F**). No changes were observed in the two control groups, i.e., mice injected with beads coupled to CD24 antibodies, but not exposed to the magnetic field (7.4 ± 1.1%, n=6 mice; **Fig. 2F)** and mice injected with beads coupled to PSA-NCAM antibodies and exposed to the magnetic field (7.4 ± 0.8%; n=6 mice; **Fig. 2F**). Importantly, the increase in Ki67+ NSCs correlated with the time the mice spent in the tunnel (**Fig. 2G**). This effect was also transient as the percentage of activated NSCs was reduced to the baseline level 21 h after exposing the mice injected with the CD24 antibody-coupled beads to the magnetic field (6.1 ± 0.5%; n=7 mice; **Fig. 2F**), which was in line with our time-lapse imaging of beads movement (**Fig. 2D**). These data indicate that constant EC cilia beating sustained the NSC quiescent state and that an acute (less than 3 h) reduction in cilia beating induced marked and transient NSC activation.

### The effect of EC cilia beating on NSC activation is mediated by Ca^2+^-permeable TRPM3

Given the marked effect of EC cilia beating on NSC activation, we next explored whether NSCs express receptors/ion channels that are either primary mechano-transducers or that are involved in mechano-sensation by amplifying the signals. To do so, we performed single nuclei RNA-sequencing (snRNA-seq) of adult mouse SVZ whole-mounts derived from three independent samples. The cell types were defined based on marker gene expression (**Fig. 3A** and **Supp. Fig. 2A**). Among the various channels/receptors that may act either as primary mechano-transducers or as mechano-amplifiers, the Ca^2+^-permeable TRPM3 showed the highest expression in the quiescent NSCs (qNSCs)/astrocyte population (**Fig. 3B**). Moreover, it was also highly enriched in qNSCs/astrocytes compared to other cell types (**Fig. 3B**). These data are consistent with a previous scRNA-seq analysis showing that the quiescent state of NSCs is characterized by a high expression of TRPM3^33^. We thus assessed TRPM3 expression at the mRNA and protein levels in adult NSCs (**Fig. 3C-G**). An RNAscope *in situ* hybridization for TRPM3, GFP, and mCherry in brain sections derived from adult mice previously electroporated with GFAP-GFP/Prom-mCherry plasmids at P0-P1 revealed that 90.2 ± 0.3% of NSCs express TRPM3 (n=195 NSCs from 3 mice; **Fig. 3D-E**). Immunolabeling for TRPM3 in a whole-mount preparation of the SVZ also showed that this receptor was expressed by GFAP-GFP+/Prom-mCherry+ NSCs that were surrounded by β-catenin+ ECs (**Fig. 3F**). The localization pattern of TRPM3 may be consistent with its presence on the apical endings/primary cilia of NSCs that were surrounded by ECs in the pinwheel structure (**Fig. 3F**). To test this possibility, we performed immunolabeling for GFP, mCherry, TRPM3 and ARL13B, the marker of primary cilia, in brain sections derived from adult mice previously electroporated with GFAP-GFP/Prom-mCherry plasmids at P0-P1 (**Fig. 3G**). This analysis revealed that TRPM3 is present in the primary cilia of NSCs (**Fig. 3G**). These data suggested that TRPM3 was enriched in qNSCs and localized at their primary cilia.

**Figure 3.**
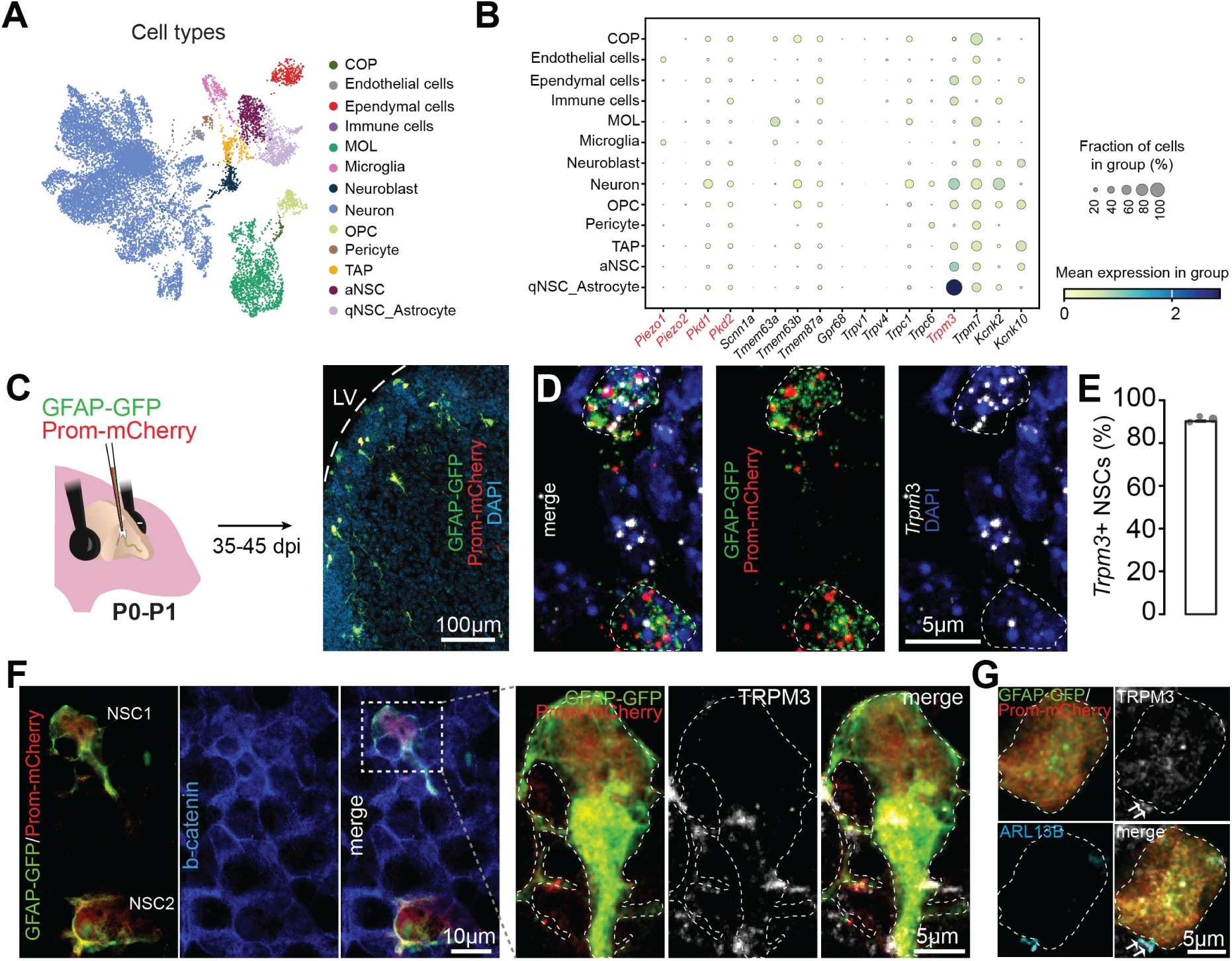
The expression of mechano-sensitive receptors in NSCs. **A:** snRNA-seq of an adult SVZ showing cell clusters defined based on marker gene expression. **B:** Dot plot showing the expression of different channels and receptors that are either directly involved in mechano-transduction or act as mechano-amplifiers in distinct cell types in the SVZ. Dot size indicates the percentage of each cell type that expresses the mRNA, and the color indicates the relative expression levels as per the key given below. The role of genes colored in red was functionally tested. **C:** Schematic drawing showing the electroporation approach used to label NSCs based on the co-expression of GFAP-GFP and Prom-mCherry and low magnification image showing NSCs. **D:** High magnification images showing an RNAscope analysis for TRPM3, GFP, and mCherry in SVZ sections derived from adult mice that had been electroporated with the GFAP-GFP and Prom-mCherry plasmids at P0-P1. **E:** Quantification of the percentage of NSCs expressing TRPM3. **F:** Immunolabeling for TRPM3, GFP, and mCherry, as well as β-catenin, an EC marker, in a whole-mount preparation of the adult SVZ. Note that GFP+ and mCherry+ NSCs were surrounded by β-catenin+ ECs in the pinwheel structure and that TRPM3 was expressed by NSCs. The white ROI outlines NSC. **G:** Immunolabeling for TRPM3, GFP, and mCherry, as well as ARL13B, a marker of primary cilia, in brain slices derived from adult mice that had been electroporated with the GFAP-GFP and Prom-mCherry plasmids at P0-P1. Note the TRPM3 expression in the ARL13B+ primary cilia of NSCs. COP, MOL, OPC and TAP indicate committed oligodendrocytes precursors, mature oligodendrocytes, oligodendrocytes precursor cells, and transient amplifying progenitors, respectively.

To determine whether TRPM3 is involved in mechano-sensation in NSCs and is required for the maintenance of their quiescent state, we used CRISPR-Cas9 to delete TRPM3 specifically in SVZ NSCs by co-electroporating a plasmid encoding a gRNA that targets *Trpm3*, Cas9, and mCherry under the Prom promoter (pU6-*Trpm3* gRNA-Prom-Cas9-T2A-mCherry) together with the GFAP-GFP plasmid (**Fig. 4A**). As a control, we used a gRNA targeting *LacZ* as previously reported^34,35^. We electroporated mice at P0-P1 and assessed NSC properties at P60 (**Fig. 4A**). We first verified whether the lack of TRPM3 affected the mechano-sensitive properties of NSCs. To do so, we measured the intracellular stiffness of NSCs as it has been previously shown that changes in the mechano-sensation and expression of different mechano-sensitive channels/receptors may lead to alterations in intracellular stiffness^36,37^. To quantify cytoplasmic stiffness at single-cell resolution in live tissue, we used a non-invasive microrheological approach^38^. This method infers intracellular mechanical compliance from spontaneous motion of endogenous organelles, without applying external forces or introducing artificial probes (**Fig. 4A-B**). We prepared acute adult brain slices derived from mice electroporated either with *TrpM3* (pU6-*Trpm3* gRNA-Prom-Cas9-T2A-mCherry) or *LacZ* (control) gRNAs, together with GFAP-GFP, incubated them with LysoTracker™ DeepRed (100 nM) for 1 h, and performed time-lapse imaging of lysosomes (**Fig. 4B**). Cytoplasmic motion was quantified by tracking individual lysosomes and computing the root mean squared displacement (RMSD) as a function of lag time (see Methods and **Supp. Fig. 2**). RMSD curves exhibited two distinct dynamical regimes, as previously described^38^. At short timescales (τ < 0.1 s), RMSD reached a high-frequency plateau corresponding to a regime in which the cytoplasm behaves as a weak elastic solid near thermodynamic equilibrium (**Supp. Fig. 2**). The plateau value (RMSD₀) provided a direct readout of cytoplasmic mechanical compliance with higher RMSD₀ values indicating a softer, more compliant intracellular environment. At longer timescales (τ > 0.1 s), RMSD increased linearly with lag time, reflecting motor-driven active transport, which was excluded from mechanical inference. Importantly, RMSD₀ fluctuations remained well above the noise floor in both conditions, ensuring reliable mechanical measurements (**Supp. Fig. 2)**. Compared to control cells, TRPM3 deletion in NSCs resulted in a significant increase in cytoplasmic mechanical compliance (Wilcoxon rank-sum test, p = 0.03), indicating cytoplasmic softening (**Fig. 4C**). These data indicated that TRPM3 was involved in mechano-sensation and that the lack of TRPM3 altered the intracellular mechanical state *in vivo*.

**Figure 4.**
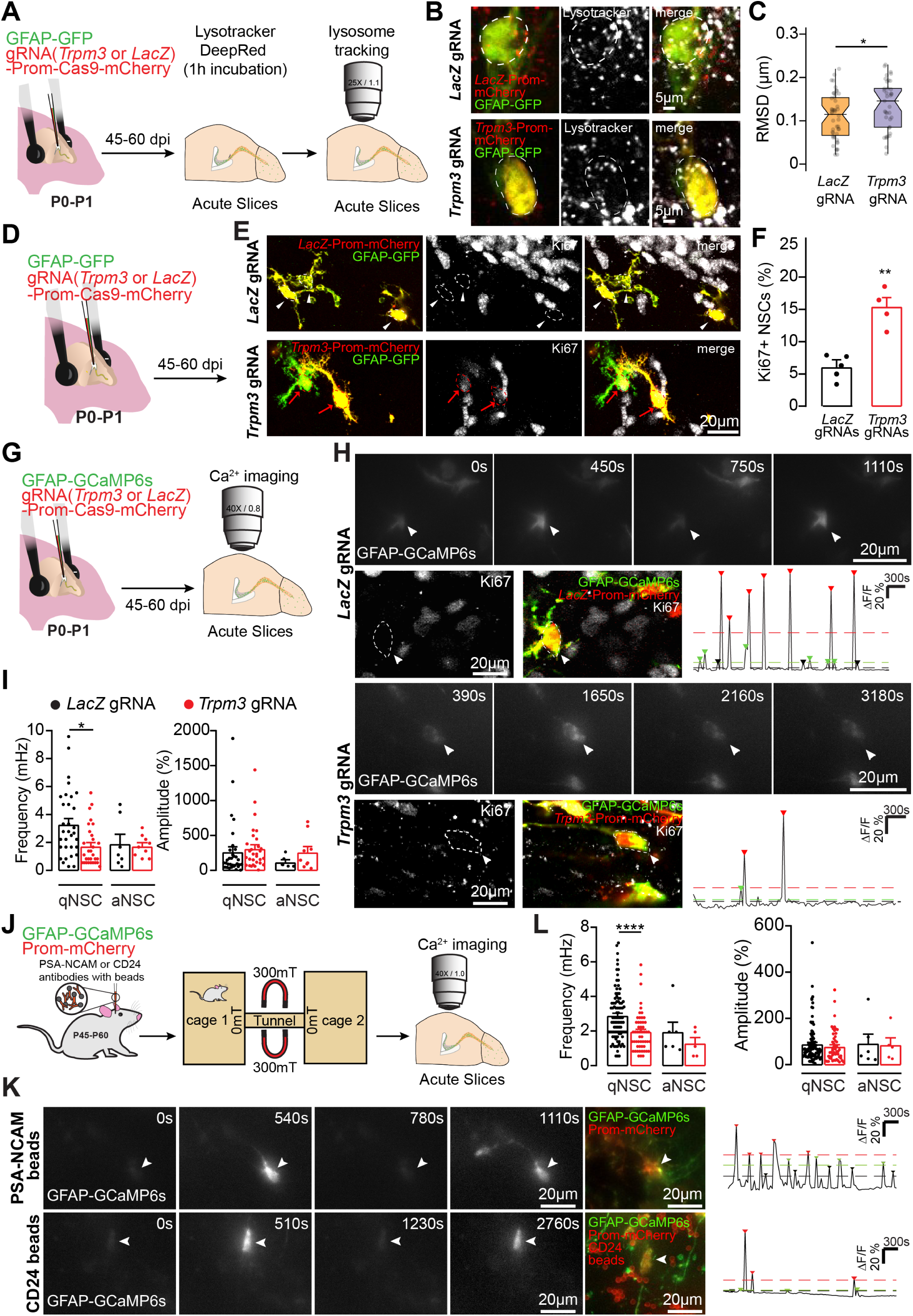
Mechano-sensitive effect of EC cilia beating on NSC activation is mediated by TRPM3. **A:** Schematic drawing of the experimental approach used for the CRISPR-Cas9-mediated deletion of *Trpm3* or *LacZ* (control) in NSCs and assessment of intracellular stiffness using lysosomes dynamics. **B:** Representative images of lysosomes (white) in NSCs with gRNAs targeting either *Trpm3* or *LacZ* (control). **C:** Assessment of intracellular stiffness of NSCs with gRNAs targeting either *Trpm3* or *LacZ* (control). **D:** Schematic drawing of the experimental approach used for the CRISPR-Cas9-mediated deletion of *Trpm3* or *LacZ* (control) in NSCs. **E-F:** Representative images and quantification of Ki67+ NSCs with gRNAs targeting either *Trpm3* or *LacZ* (control). The red arrows and ROI indicate Ki67+ NSCs while the white arrowheads and ROI indicate Ki67- NSCs. **G:** Schematic drawing of the experimental approach used for the CRISPR-Cas9-mediated deletion of *Trpm3* in NSCs and the assessment of Ca^2+^ dynamics. We co-electroporated gRNAs against *Trpm3* or *LacZ* (control) together with Prom-Cas9-mCherry and GCaMP6s, a Ca^2+^ sensor, under the GFAP promoter, at P0-P1 and prepared acute brain slices from 2-3-month-old mice. **H:** Representative images and traces of Ca^2+^ activity in NSCs with gRNAs targeting either *Trpm3* or *LacZ* (control). The white arrowheads and ROIs indicate Ki67- qNSCs. **I**: Quantification of the frequency and amplitude of Ca^2+^ events in qNSCs and aNSCs under control *LacZ* conditions and following *Trpm3* deletion. **J:** Schematic drawing of the experimental approach used for the assessment of Ca^2+^ activity in NSCs following EC cilia inhibition *in vivo*. The adult mice, which were previously electroporated with GFAP-GCaMP6s and Prom-mCherry at P1, were placed in the behavioral cages connected by a tunnel equipped with permanent magnets for 3 h, following the injection of the CD24- or PSA-NCAM antibody-coupled magnetic beads. The acute brain slices were prepared immediately after and Ca^2+^ activity was assessed in the areas containing the magnetic beads coupled to EC cilia. **K:** Representative images and traces of Ca^2+^ activity in NSCs following the inhibition of EC cilia beating. **L**: Quantification of the frequency and amplitude of Ca^2+^ events in qNSCs and aNSCs under control condition (PSA-NCAM antibody-coupled beads+magnet) and following the inhibition of EC cilia beating (CD24 antibody-coupled beads+magnet). Statistical significance was determined using an unpaired Student’s t-test. \**p*<0.05 and \*\**p*<0.01. Data are expressed as means ± SEM. N=22 and 23 NSCs from 4 mice per group for LacZ and Trpm3 gRNAs, respectively, for **C,** n=353 and 551 NSCs from 5 and 4 mice for LacZ and Trpm3 gRNAs, respectively, for **F**, n=32 and 30 NSCs from 9 and 6 mice, respectively, for qNSCs and 9 and 7 NSCs from 4 mice for aNSCs, respectively for **I**, and n=95 and 53 qNSCs and n=6 and 5 aNSCs from 3 PSA-NCAM antibody-coupled beads + magnet and 5 CD24 antibody-coupled beads + magnet mice, respectively, for **L**.

We next determined whether the lack of TRPM3 would affect NSC activation by inducing their proliferation (**Fig. 4D**). Deleting *Trpm3* in NSCs induced a marked increase in the number of proliferative NSCs (6.1 ± 0.8% vs. 15.2 ± 1.5% in mice electroporated with *LacZ* and *Trpm3* gRNAs, respectively; **Fig. 4E-F**). Together with the assessment of activated NSCs after abrogated EC cilia beating, these data indicated that both, the lack of TRPM3 receptor in NSCs (**Fig. 4F**) and the abrogation of mechano-sensitive stimuli by inhibiting EC cilia beating (**Fig. 2F-G**) induced marked NSC activation, suggesting that constant EC cilia beating may be required to maintain the quiescent state of NSCs involving TRPM3.

TRPM3 is a Ca^2+^ permeable receptor^39,40^ and our previous observations highlighted the important role that Ca^2+^ activity plays in the regulation of NSC quiescence/activation dynamics^34,35^. We thus determined whether a lack of TRPM3 in NSCs would affect the frequency and amplitude of Ca^2+^ events in NSCs in their quiescent (qNSCs) and activated (aNSCs) states. To do so, we electroporated the Ca^2+^ sensor GCaMP6s under the GFAP promoter together with gRNAs targeting either *LacZ* or *Trpm3* as well as Cas9 and mCherry under the Prom promoter at P0-P1 and performed Ca^2+^ imaging in acute brain sections at P45-P60 (**Fig. 4G**). After 1 h of imaging, we fixed brain sections and performed *post hoc* immunolabeling for Ki67 to determine whether imaged NSCs were in the quiescent or activated state (**Fig. 4H**). Interestingly, the lack of *Trpm3* in NSCs resulted in a decrease in the frequency of Ca^2+^ fluctuations in qNSCs but not in aNSCs (**Fig. 4H-I**). No difference in the amplitude of Ca^2+^ events was observed after *Trpm3* deletion, although a tendency, albeit not significant, to an increased amplitude was seen in aNSCs (**Fig. 4I**). We then determined whether the same changes in the Ca^2+^ activity could be also observed after inhibiting EC cilia beating. To do so, we injected either CD24- or PSA-NCAM antibody-coupled magnetic beads into the LV of adult mice previously electroporated with the GFAP-GCaMP6s and Prom-mCherry plasmids at P1 and placed the mice in the behavioral cages connected by a tunnel equipped with permanent magnets for 3 h (**Fig. 4J**). Immediately after the 3-h period we prepared acute brain slices and performed time-lapse imaging of Ca^2+^ responses in NSCs in the regions surrounded by magnetic beads coupled to EC cilia (**Fig. 4K**). Our analysis revealed that there was a marked decrease in the frequency but not the amplitude of Ca^2+^ events in qNSCs (**Fig. 4K-L**). These data are consistent with the effect observed after TRPM3 deletion and suggest that a decreased frequency of Ca^2+^ events in qNSCs might lead to their activation, as we reported previously^34,35^.

### PKD1/2 are expressed in NSCs and act as mechano-transducers

Although TRPM3 may mediate mechano-sensitive responses in various cells by amplifying the signal, it is not thought to be a primary mechano-transducer^41^. We thus explored the function of primary mechano-transducers such as Piezo1/2 and PKD1/2 in NSCs. Our snRNA-seq analysis revealed that Piezo1/2 were not expressed in adult NSCs (**Fig. 3B**). Consistent with the expression analysis, CRISPR-Cas9 mediated deletion of both Piezo1 and Piezo 2 in adult NSCs did not affect NSCs proliferation (**Fig. 5A-C**). Similarly, bath application of Yoda 1, a Piezo 1 activator, did not affect Ca^2+^ activity in NSCs (**Supp. Fig. 2**). Thus, our snRNA-seq, CRISPR-Cas9 mediated deletion of Piezo1/2 and pharmacological activation experiments suggested that Piezo1/2 were not involved in the effect mediated by the inhibition of EC cilia beating.

**Figure 5.**
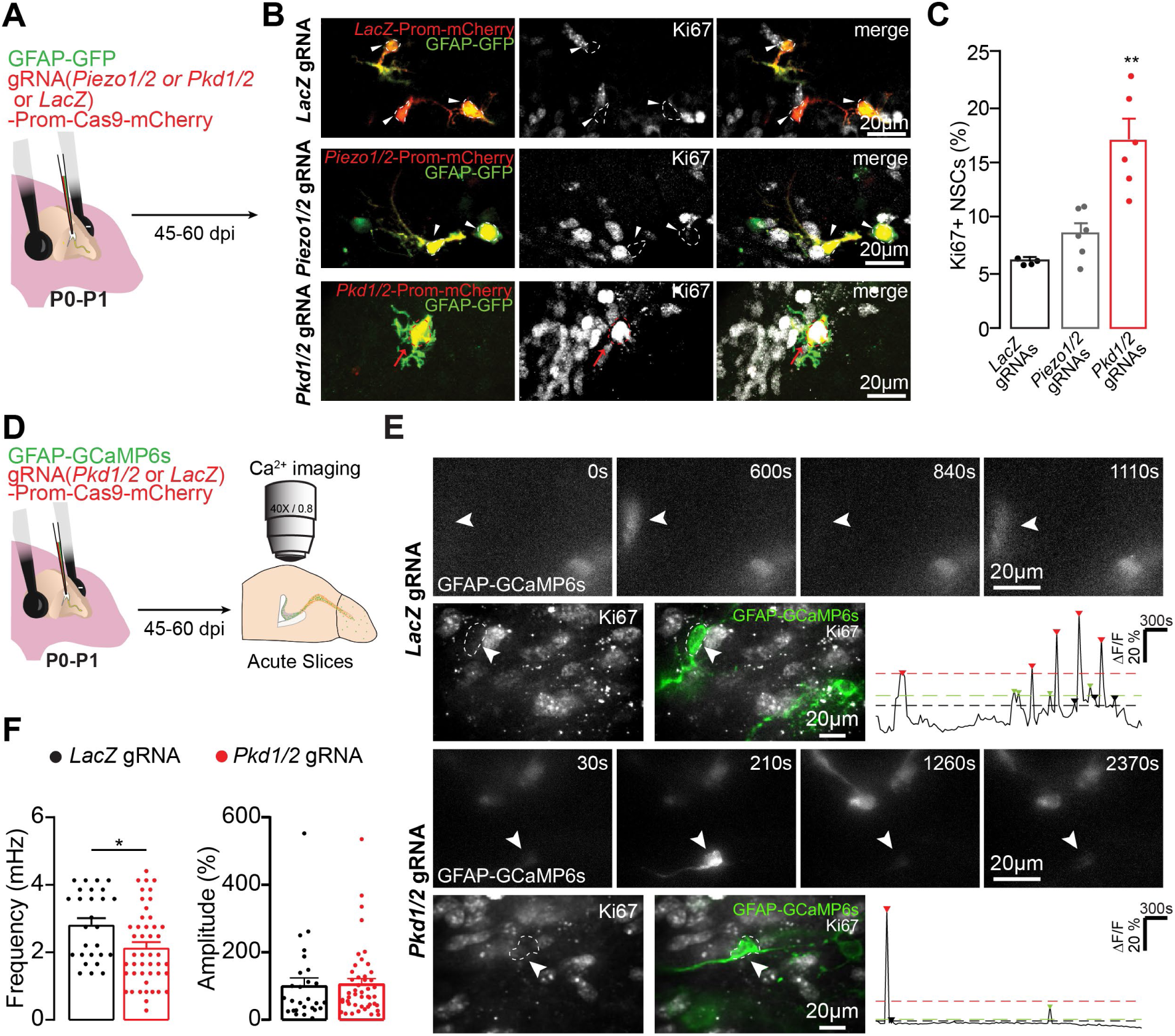
PKD1/2, but not Piezo1/2, regulate the effect of EC cilia beating on NSCs. **A:** Schematic drawing showing the electroporation approach used for the CRISPR-Cas9-mediated deletion of *Piezo1/2 or Pkd1/2* (or *LacZ* as a control) in NSCs. **B-C:** Representative images and quantification of Ki67+ NSCs with gRNAs targeting either *LacZ* (control) or *Piezo1/2* or *Pkd1/2*. The red arrows and ROI indicate Ki67+ NSCs while the white arrowheads and ROI indicate Ki67- NSCs. **D:** Schematic drawing of the experimental approach used for the CRISPR-Cas9-mediated deletion of *Pkd1/2* in NSCs and assessment of Ca^2+^ dynamics. **E:** Representative images and traces of Ca^2+^ activity in NSCs with gRNAs targeting either *LacZ* or *Pkd1/2*. The white arrowheads and ROIs indicate Ki67- qNSCs. **F**: Quantification of the frequency and amplitude of Ca^2+^ events in NSCs under control *LacZ* conditions and following *Pkd1/2* deletion. Statistical significance was determined using an unpaired Student’s t-test. \**p*<0.05 and \*\**p*<0.01. Data are expressed as means ± SEM. N=511, 347 and 388 NSCs from 4, 6 and 6 mice for *LacZ, Piezo1/2*, and *Pkd1/2* gRNAs, respectively, for **C;** 46 and 27 qNSCs for *Pkd1/2* and *LacZ* gRNAs, respectively, for **E**.

We next focused on PKD1/2, which are expressed at higher levels in adult NSCs compared to Piezo1/2 (**Fig. 3B**). It has been also previously shown that PKDs are expressed in immotile cilia and mechanically sense the fluid flow required for left-right determination during embryonic development^42,43^. We prepared gRNAs targeting both *Pkd1* and *Pkd2* (pU6-*Pkd1* gRNA-Prom-Cas9-T2A-mCherry and pU6-*Pkd2* gRNA-Prom-Cas9-T2A-mCherry), or *LacZ* (control), and electroporated them together with GFAP-GFP into P1 pups (**Fig. 5A**). We analyzed the percentage of activated Ki67+ NSCs in the adult SVZ and observed a marked increase in their proliferation (**Fig. 5B-C**). As PKDs are Ca^2+^-permeable channels^41^, we also determined whether the lack of PKD1/2 affected Ca^2+^ dynamics in NSCs. To do so, we electroporated gRNAs targeting both *Pkd1* and *Pkd2* (pU6-*Pkd1* gRNA-Prom-Cas9-T2A-mCherry and pU6-*Pkd2* gRNA-Prom-Cas9-T2A-mCherry), or *LacZ* (control), together with GFAP-GCaMP6s into P1 pups and assessed Ca^2+^ activity in NSCs in acute brain slices derived from adult mice (**Fig. 5D**). Our analysis revealed that there was a decrease in the frequency but not the amplitude of Ca^2+^ events in NSCs targeted with *Pkd1/2* gRNAs compared to control, *LacZ*-expressing gRNAs (**Fig. 5E-F**). These data were consistent with our analysis of TRPM3-mediated effects following EC cilia inhibition and suggested that PKD1/2 and TRPM3 might act in concert to mediate mechanical stimuli-induced NSC quiescence, with PKD1/2 being the primary mechano-transducer in NSCs, and with TRPM3, which is highly enriched in NSCs, amplifying the signal.

### EC cilia beating inhibition triggers NSCs activation through increased protein translation

We next explored the mechanisms by which PKD1/2-TRPM3 mediated effects of EC cilia beating inhibition triggered NSCs proliferation. To do so, we performed an snRNA-seq of the SVZ after inhibiting EC cilia beating (**Fig. 6A**) and compared it to the dataset derived from control samples (**Fig. 3A** and **Fig. 6B**). No difference was observed in the distribution of clusters between control samples injected with CD24 antibody-coupled beads and not exposed to the magnets compared to those that were injected with CD24 antibody-coupled beads and were exposed to the magnets (**Fig. 6B**). Our analysis also did not reveal any changes in the expression of mechano-receptors after EC cilia beating inhibition (**Fig. 6C**). To explore how EC cilia inhibition triggers NSC activation, we performed an enrichment analysis for Gene Ontology (GO) Biological Process (BP) terms for the genes significantly upregulated in aNSCs after inhibiting cilia beating of EC (**Fig. 6D**). Our analysis revealed significant changes in the GO terms “negative regulators of ryanodine-sensitive Ca^2+^ release channel activity”, “regulators of ryanodine-sensitive Ca^2+^ release channel activity” “negative regulators of Ca^2+^ ion export across plasma membrane”, “detection of Ca^2+^ ions” and several other GO terms related to Ca^2+^ signaling (**Fig. 6D**). These data are in line with our Ca^2+^ imaging analysis and suggest that alterations in the Ca^2+^ fluctuations in NSCs induced by inhibited EC cilia beating could induce Ca^2+^-dependent transcriptional program to trigger proliferation. We have also observed a marked changes in GO term “cytoplasmic translation” (**Fig. 6D**), which is also known to be involved during transitions from quiescence to activation and early proliferation^44,45^. Particularly important were changes in genes involved in Ca^2+^ signaling such as *Calm1, Calm2, Calm3, Ghitm, Ptprj*, and protein translation including *Rplp1, Rpl26, Eif5a, Rpl13, Rps11, Rps25, Rpl19, Rps3a1, Rps29, Rps21, Rpl18a, Rpl3, Rps7 and Fau* (**Fig. 6E**).

**Figure 6.**
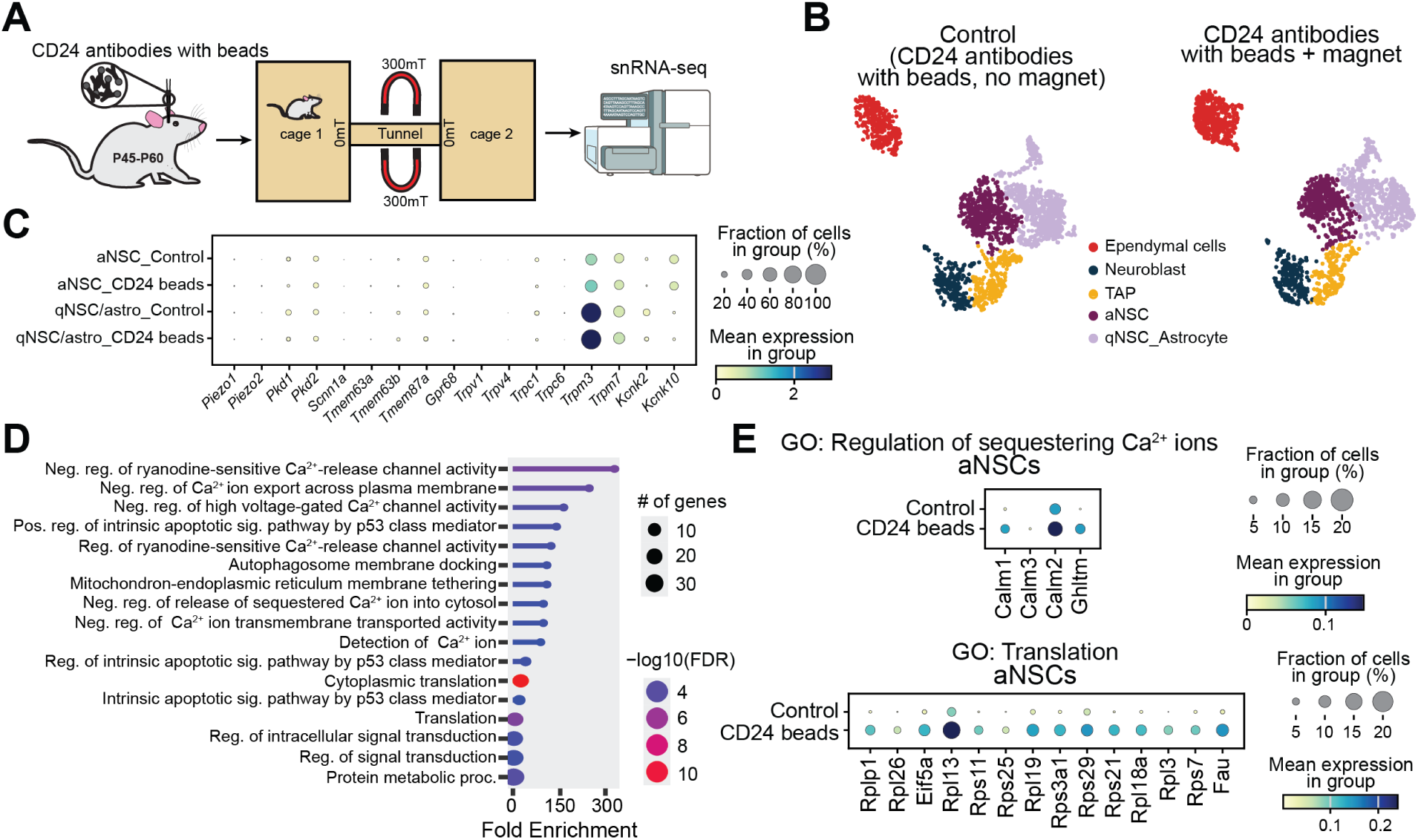
Inhibition of EC cilia beating triggers NSCs activation through increased protein translation. **A:** Schematic drawing of the experimental approach used for the modulation of EC cilia beating by a magnetic field and snRNA-seq analysis. **B**: Cell types defined based on the marker gene expression of an snRNA-seq dataset of adult SVZ derived from mice injected with CD24 antibody-coupled beads and placed in the behavioral cages connected by a tunnel that was either equipped (CD24 beads + magnet) or lacked (control) permanent magnets. **C:** Dot plot showing the expression of different mechano-sensitive receptors in NSCs in the control condition and following EC cilia inhibition. The dot size indicates the percentage of each cell type that expressed the mRNA, and the color indicates the relative expression levels. Note that there are no differences in the expression patterns. **D:** GO enrichment analysis in aNSCs for biological process terms following the inhibition of EC cilia beating. **E:** Dot plot showing the expression of different genes either involved in Ca^2+^ signaling or protein translation in aNSCs under control conditions and following the inhibition of EC cilia beating. The dot size indicates the percentage of each cell type that expressed the mRNA, and the color indicates the relative expression levels.

### *In vivo* pharmacological activation of TRPM3 rescues NSC activation induced by abrogated EC cilia beating

The results reported above showed that the deletion of *Trpm3* and *Pkd1/2* in NSCs or the inhibition of EC cilia beating for a maximum of 3 h resulted in marked NSC activation through increase in cytoplasmic translation. Although these data are consistent with a mechano-sensitive influence on NSC activation by EC cilia beating, it is possible that the 3-h arrest of cilia beating also affects the presence of various molecular cues in the cerebrospinal fluid that in turn trigger NSC activation independently of TRPM3. To address this possibility, we determined whether the effects mediated by the inhibition of EC cilia beating and the lack of TRPM3 were additive or inhibiting each other. We reasoned that if the effects induced by the inhibition of EC cilia beating were independent of TRPM3 then inhibiting EC cilia beating would further increase the proliferation of NSCs that lack *Trpm3*. However, if EC cilia beating-induced effects on NSC activation depended on TRPM3, the lack of TRPM3 would impede any additional NSC activation induced by EC cilia inhibition. We therefore blocked EC cilia beating for a maximum of 3 h (**Fig. 7A**), as described above, but in mice that had been electroporated either with gRNAs against *Trpm3* or *LacZ* (control) as well as Cas9 and mCherry under the Prom promoter together with GFAP-GFP at P0-P1 (**Fig. 7A**). At P45-P60, the mice were injected in the LV with beads coupled to CD24 antibodies and were placed in the behavioral cages connected by a tunnel equipped with permanent magnets for 3 h (**Fig. 7A**). Our data revealed that the deletion of *Trpm3*, that by its own increases NSC proliferation (**Fig. 4F**), impeded additional NSC activation induced by the altered EC cilia beating (17.9 ± 2.2% for mice electroporated with *LacZ* gRNA and injected with CD24 antibody-coupled beads vs. 16.5 ± 3.1% in mice electroporated with *Trpm3* gRNAs and injected with CD24 antibody-coupled beads; **Fig. 7B-C**). These data indicate that EC cilia beating had a mechano-sensitive effect on NSC quiescence/activation dynamics through TRPM3-dependent pathways.

**Figure 7.**
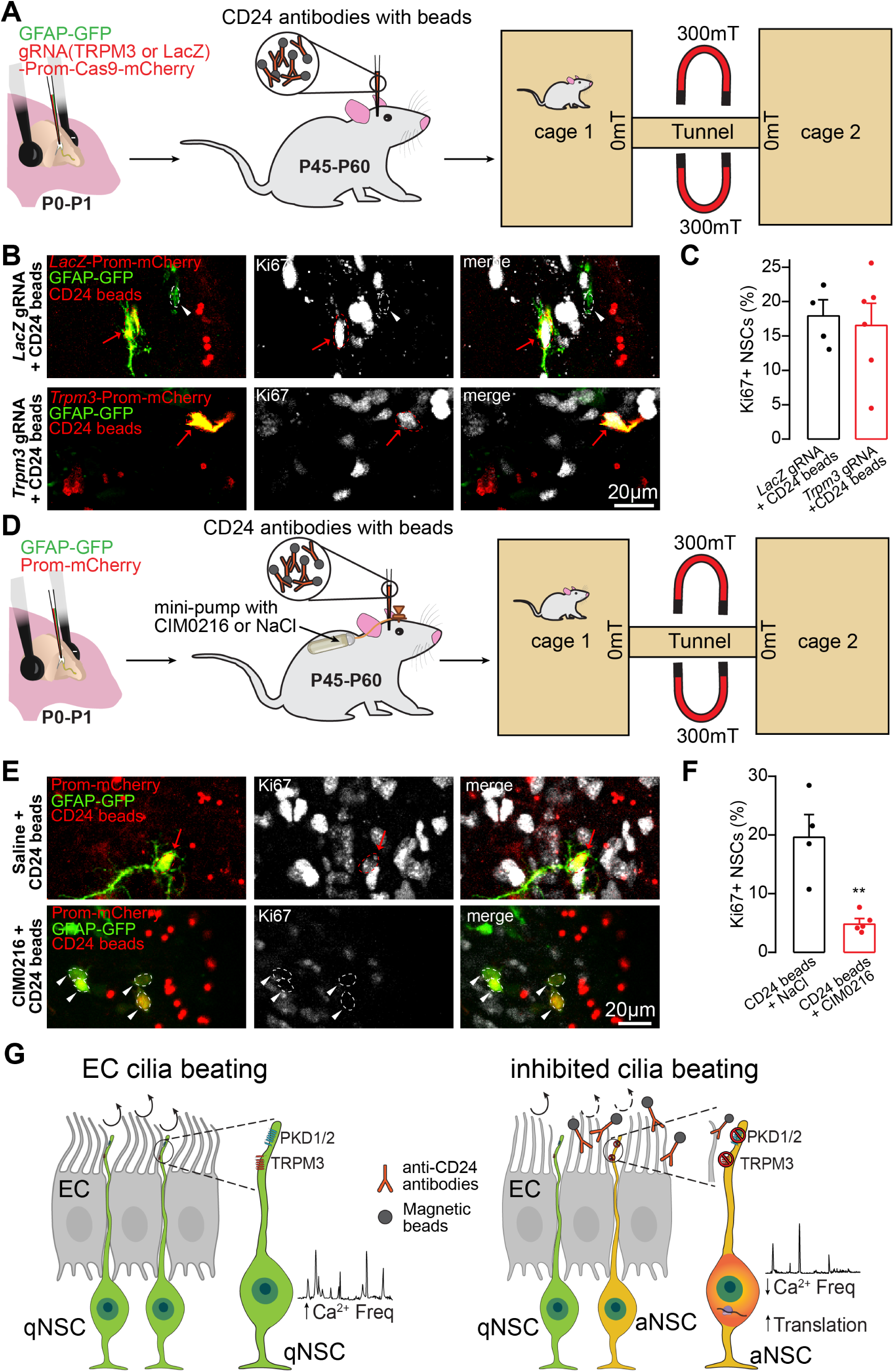
*In vivo* activation of TRPM3 reverses the NSC transition into the activated state following decreased EC cilia beating. **A:** Schematic drawing of the experimental approach used for the CRISPR-Cas9-mediated deletion of *Trpm3* (or *LacZ* as a control) in NSCs and modulation of EC cilia beating by a magnetic field. Following the injection of CD24 antibody-coupled beads, the mice were placed for 3 h in a behavioral chamber consisting of two cages connected by a tunnel equipped with permanent neodymium magnets. **B-C:** Representative images and quantification of Ki67+ NSCs with gRNAs targeting either *LacZ* (control) or *Trpm3* after the arrest of EC cilia beating for 3 h. **D:** Experimental approach used for the pharmacological activation of TRPM3 and the modulation of EC cilia beating by a magnetic field. Osmotic minipumps containing either NaCl or CIM0216, a selective TRPM3 agonist, were installed during the same surgery as for the injection of the CD24 antibody-coupled beads. The mice were placed for 3 h in the behavioral cages connected with a tunnel equipped with permanent neodymium magnets. **E-F:** Representative images and quantification of Ki67+ NSCs following the infusion of either NaCl or CIM0216 while blocking EC cilia beating for a maximum of 3 h. Note that the activation of TRPM3 alleviated EC cilia beating inhibition-induced NSC activation. Statistical significance was determined using an unpaired Student t-test. \*\**p*<0.01. Data are expressed as means ± SEM. N=223 and 283 NSCs from 4 and 6 mice, respectively, for **C**, and n=286 and 399 NSCs from 4 and 5 mice installed with osmotic minipumps containing either NaCl or CIM0216, respectively, for **F**. The white arrowheads and ROIs indicate Ki67- qNSCs while the red arrows and ROIs indicate Ki67+ aNSCs. **G:** Schematic showing the interplay between EC cilia beating and NSC quiescence/activation dynamics through mechano-sensation and downstream Ca^2+^ activity and protein synthesis.

We next determined whether the pharmacological activation of TRPM3 would alleviate the effects induced by the reduction of ECs cilia beating frequency. To test this possibility, during the same surgery used to inject beads coupled to CD24 antibodies into the LV, we installed an osmotic minipump filled with CIM0216, a selective TRPM3 agonist. The osmotic mini-pump was connected to a plastic-based cannula and infused the CIM0216 into the LV. In control mice, the osmotic mini-pump was filled with 0.9% saline. After a 30-45 min recovery period, the mice were allowed to freely explore the behavioral cages connected by a tunnel equipped with permanent magnets for 3 h (**Fig. 7D**). Interestingly, the activation of TRPM3 in NSCs while blocking ECs cilia beating reversed NSC activation and maintained NSCs in their quiescent state (19.8 ± 3.7% for mice infused with NaCl and injected with CD24 antibody-coupled beads vs. 5.1 ± 0.7% for mice infused with CIM0216 and injected with CD24 antibody-coupled beads; **Fig. 7E-F**).

Altogether our data revealed that constant ECs cilia beating created mechanical forces that, through PKD1/2-TRPM3 mechano-sensation in NSCs and downstream Ca^2+^ signaling, maintained NSCs in the quiescent state (**Fig. 7G**).

## Discussion

Here, we revealed a dynamic interplay between ECs and NSCs, pinpointing on the source of mechanical forces created by EC cilia beating that enforce NSC quiescence. We showed that EC cilia beating sustains the quiescent state of NSCs through PKD1/2-TRPM3 pathway and downstream Ca^2+^ signaling. As a high frequency of Ca^2+^ fluctuations is required to maintain the quiescent state of NSCs^34,35,46^, we suggest that changes in the Ca^2+^ entry through PKD1/2-TRPM3 mechanistically link EC cilia beating to NSC quiescence/activation dynamics. An acute (less than 3 h) abrogation of EC cilia beating or lack of PKD1/2 or TRPM3 in NSCs alters Ca^2+^ dynamics and leads to NSC activation linked to Ca^2+^ signaling and increased protein translation.

Although previous studies have demonstrated the mechano-sensitive properties of NSCs by showing that NSC activation and differentiation are regulated by the stiffness of the substrates *in vitro*^22,24^ or shear stress *ex vivo*^23^, the potential source of the mechanical forces generated *in vivo* was unclear. Our data suggest that the constant beating of ECs’ cilia that surround NSCs in a pinwheel structure generate mechanical forces that retain NSCs in quiescence as decreased cilia beating triggers NSC activation. Our work also showed that PKD1/2 and TRPM3, but not Piezo1/2, controls adult NSC quiescence/activation dynamics. This is in line with previous work demonstrating the role of other mechano-sensitive receptors such as the epithelial sodium channel (ENaC) in the regulation of adult NSCs^23^. Interestingly, however, while the lack of ENaC, which mediates shear stress responses in adult NSCs and their progeny impairs progenitor proliferation and neurogenic output in the olfactory bulb^23^, the lack of TRPM3 or PKD1/2, in contrast, results in NSC activation. Since sheer stress was applied to the entire surface of an SVZ whole-mount preparation^23^, while here we specifically blocked ECs cilia beating, it would be interesting to explore if and how shear stress affects EC cilia beating. It has been previously reported that EC cilia beating withstands the forces produced by the external fluid flow^47^, indicating that shear stress may have affected progenitor proliferation through the mechanisms independent on EC cilia beating. This in turn suggest that impact mechanical forces on NSCs may be context dependent. Our data with PKD1/2 are also in line with previous reports showing that the presence of these mechano-sensitive receptors in embryonic NSCs/radial glia is required for the planar polarization of the brain ventricular epithelium, while their lack affects the development and function of NSCs and EC^48^. Furthermore, it has been demonstrated that PKD2 is preferentially localized on immotile cilia at the ventral node of mouse embryos and is required for sensing fluid flows for the left-right symmetry of the body^42,43^. This is reminiscent of the presence of PKD1/2 in adult NSCs that bear immotile primary cilia and sense mechanical forces generated by the beating of EC cilia. Interestingly, among all the primary mechano-transducers and channels/receptors involved in mechano-sensation, the expression of TRPM3 was highest. Although TRPM3 is not thought to be a primary mechano-transducer, it may be involved in amplifying the signals generated by PKD1/2, as other TRP channels do^41^. Further studies are required to elucidate the exact molecular interaction between PKD1/2 and TRPM3 channels in NSCs.

Our data also revealed that the mechano-sensitive effects on NSCs are mediated by Ca^2+^ activity, which is in line with our previous reports showing a major role in Ca^2+^ activity in NSCs for integrating and decoding various niche-derived signals that determine the activation or quiescence dynamics of NSCs^23,34,35^. Interestingly, our snRNA-seq data revealed that the inhibition of EC cilia beating was linked to changes in Ca^2+^ signaling and increased protein translation. These processes can be also linked. Ca^2+^ is one of context-dependent regulators of protein translation. For example, it has been demonstrated that the pharmacological blockade of Ca²⁺ entry into neurons increases translation rates in neurons^49^. Similarly, in stem/progenitor cells, Ca^2+^ influx leads to the inhibition of mTORC1 signaling^50^, thereby suppressing translation. This is line with our data showing that EC cilia beating-generated Ca^2+^ fluxes in NSCs were associated with reduced protein translation, while the inhibition of cilia beating, which results in decreased Ca^2+^ fluxes into NSCs, increases protein translation and triggers NSCs proliferation.

### Limitations of the study

In the present work we focused on Ca^2+^ imaging at the cell soma level of NSCs. It would be interesting to decipher Ca^2+^ dynamics in the apical processes and primary cilia of NSCs where TRPM3 and PKD1/2 are localized and strategically positioned to mediate EC cilia beating-driven responses. It should be also noted that while more than 85% of electroporated cells are NSCs^35^, we cannot formally rule out that CRISPR-Cas9 deletion of TRPM3 or PKD1/2 could also have affected EC, in addition to NSCs, since Cas9 was under the Prom promoter. However, TRPM3 or PKD1/2 are expressed at much lower level in other cell types compared to NSCs.

Lastly, cilia formation and beating are modulated by several physiological and pathological conditions, such as circadian rhythm, hyperglycemia, alcohol consumption and various neurodevelopmental and neurodegenerative disorders^25-29,51^. Since these conditions can be also linked to changes in the progenitors’ proliferation and brain development, it would be interesting to explore how changes in cilia formation and function affect NSCs physiology under these conditions. It remains also unclear whether ECs’ cilia beating arrest-induced activation of NSCs results in a neurogenic or gliogenic differentiation potential and whether, over the long-term, it may result in alterations in neural circuitry formation. This in turn may shed light on the underlying mechanisms of some of neurodevelopmental disorders induced by alterations in the formation/function of ECs cilia^2,11,20,52^.

Altogether, our data revealed an exquisite interplay between ECs’ cilia beating and NSC quiescence and activation dynamics. They further pointed to the sources of mechanical forces that may influence NSCs and to some key mechano-sensitive mechanisms operating in NSCs to regulate their quiescence and activation.

## RESOURCE AVAILABILITY

### Lead Contact

Further information and requests for resources and reagents should be directed to and will be fulfilled by the Lead Contact, Armen Saghatelyan.

### Material Availability

This study did not generate new unique reagents.

### Data and code Availability

The code for the analysis of Ca^2+^ activity in NSCs is available at https://github.com/SagLab-CERVO/Calcium_analysis_in_NSC. GEO accession number for snRNAseq datasets under control condition (GSM9518782, GSM9518783, GSM9518784) and following EC cilia inhibition (GSM9518785, GSM9518786, GSM9518787) is # GSE319498. The datasets can be accessed at GEO accession GSE319498 (https://www.ncbi.nlm.nih.gov/geo/query/acc.cgi?acc=GSE319498).

## Acknowledgments

We thank the Cell Biology and Image Acquisition (CBIA) Core Facility (RRID:SCR_021845) at the University of Ottawa for the use of the various imaging systems and Marie-Anne Lebel-Cormier for optimizing the MATLAB script for the bead movement analysis. This work was supported by a Canadian Institute of Health Research (CIHR) grant to A.S. A.S. is a Canada Research Chair (Tier I) in postnatal neurogenesis.

## Author contributions

C.B., A.G. and R.R-A performed the electroporations and cilia movement analyses *ex vivo*. C.B. and R.R-A performed all the *in vivo* experiments. M.S. developed the CRISPR-Cas9 model in the NSCs and performed the RNAscope *in situ* hybridizations. C.B., M.S., and A.C. performed the confocal imaging and analyses. N.R. and J.B. generated snRNA-seq dataset. M.L.R. and J. F-S analyzed snRNA-seq data. M.G. supervised snRNAseq analysis and revised the manuscript. E.H. assessed intracellular stiffness of NSCs. A.S. supervised the project and wrote the manuscript, considering the comments of all the authors.

## Material and Methods

### Animals

The experiments were performed using one- to three-month-old CD1 (Charles River) mice of both sexes, which were electroporated on postnatal days 0-1 (P0-P1). All experiments were approved by the Université Laval and University of Ottawa animal protection committees. The mice were kept on a 12-h light/dark cycle at a constant temperature (22°C) with food and water *ad libitum*.

### Electroporation

For the electroporation of plasmids *in vivo*, P0-P1 CD-1 pups were anesthetized by hypothermia and were placed on a stereotactic frame. The plasmids (1 µL, 3-6 µg/µL total, except for GFAP-GCaMP6s) were injected into the lateral ventricle (LV) using the following coordinates (with respect to the lambda): anterior-posterior (AP) 1.8 mm, medio-lateral (ML) 0.8 mm, and dorso-ventral (DV) 1.6 mm. For GFAP-GCaMP6s, no more than 1 µg/µL of 1 µL of solution was used to avoid buffering the intracellular Ca^2+^ by GCaMP6s, a Ca^2+^ indicator. We used the following plasmids: CMV-GCaMP6s (#40753, Addgene), kindly provided by Dr. Kim & GENIE Project, Janelia Research Campus^53^; CMV-GFP (#U55763, Clontech), PU6-BbsI-CBh-Cas9-T2A-mCherry (#64324, Addgene), kindly provided by Dr. Kuehn, Berlin Institute of Health^54^, and CMV-Lck-GFP (#61099, Addgene), kindly provided by Dr. Green, University of Iowa^55^. GCaMP6s and GFP were sub-cloned under the GFAP promoter, while mCherry derived from the PU6-BbsI-CBh-Cas9-T2A-mCherry plasmid was subcloned under the prominin (Prom) promoter, as described previously^32,34,35^. The plasmid solutions were supplemented with 1% Fast-Green dye to visualize the injection and ensure that the ventricle was filled with plasmid solution. Following the injection, an electric field (five 50-ms 100-mV pulses at 950-ms intervals) was quickly applied using an electrode coated with conductive gel placed on the bone surface of the pup. Electroporated animals were used at P45-P90.

For the electroporation of CMV-Lck-GFP, P30-P45 mice were used, which resulted in better labeling of ependymal cells (ECs). The following coordinates were used for CMV-Lck-GFP injections into the LV (with respect to the bregma): AP 0.3 mm, ML 0.9 mm, and DV 2.3 mm, and an electric field of five 50-ms 150-mV pulses at 950-ms intervals was quickly applied. The animals were used 5-7 days post-electroporation.

#### CRISPR target site selection and assembly

To perform CRISPR-Cas9 editing in NSCs, we used *Trpm3* gRNAs designed and selected using ChopChop online software^56^. Validated gRNAs were used to target the *LacZ* gene as a control, as previously reported^34,35^. Two different gRNAs were used to target *Trpm3* to increase the efficiency of the CRISPR editing and were electroporated together. For the Piezo1/2 experiments, one gRNA was used for Piezo1 and another for Piezo2. Both gRNAs were mixed and injected at the same time together with the GFAP-GFP plasmid into the LV. We used the same approach for PKD1/2. The following gRNA sequences were used:

gRNA1-Trpm3 GGTTTTCCACAATTCCCCAG

gRNA2-Trpm3 GGAGTGCATGCTATTGAGAA

gRNA-Piezo1 TCCTCGTGAGGCGTCCACAG

gRNA-Piezo2 GCAATGAACATCCCGATATC

gRNA-PKD1 TGGAATGGGCCCCCTGCACG

gRNA-PKD2 TAAAAGCTGCCCGCGTGCCC

The gRNAs were cloned in the BbsI site of the PU6-BbsI-CBh-Cas9-T2A-mCherry plasmid in which CBh was replaced by the Prom promoter.

#### Immunostaining

The animals were deeply anesthetized with sodium pentobarbital (12 mg/mL; 0.1 mL per 10 g of body weight) and were perfused intracardially with 0.9% NaCl followed by 4% paraformaldehyde (PFA). The brains were collected and were kept overnight in 4% PFA. Sagittal slices (40 µm) were cut using a vibratome (Leica). In some experiments, immunolabeling was performed in whole-mount preparations of the SVZ that had been dissected as described by Mirzadeh et al.^57^. The brain slices or SVZ whole-mounts were incubated with the following primary antibodies: chicken anti-GFP (RRID: AB_10000240; 1:1000, Avès), rat anti-mCherry (RRID: AB_2536611; 1:1000, Thermo Fisher Scientific), rabbit (RRID: AB_443209; 1:1000, Abcam) or mouse (RRID: AB_2797703; 1:500, Cell Signaling) anti-Ki67, rabbit anti-TRPM3 (RRID:AB_1291859; 1:500, Novus Biologicals), mouse anti-Arl13B (RRID:AB_3073658; 1:500, Abcam), rabbit anti-Rpl13 (RRID:AB_1856433; 1:1000, Sigma), rabbit anti-Rps11 (RRID:AB_3268408; 1:1000, Novus) and mouse anti-β-catenin (RRID:AB_397554; 1:500, BD Bioscience). The antibodies were diluted in 0.5% Triton X-100 and 4% milk prepared in PBS. The brain slices were incubated overnight at 4°C. The corresponding secondary antibodies were used and were applied for 3 h at RT. Fluorescence images were acquired either using a confocal microscope (FV 1000; Olympus) with 60x (UPlanSApoN 60x/NA 1.42; Olympus) and 40x (UPlanSApoN 40x/NA 0.90; Olympus) oil and air immersion objectives, respectively, or using a confocal microscope (LSM 700; Zeiss) with a 20x air immersion objective (EC Plan-Neofluar 20x/NA 0.50; Zeiss) for low magnification images or a 40x oil immersion objective (Plan-APOCHROMAT 40x/NA 1.4; Zeiss).

#### Coupling of CD24, PSA-NCAM and PDGFRα antibodies to the magnetic beads

Rat CD24 (RRID: AB_312834, Biolegend) or mouse PSA-NCAM (RRID: AB_95211, MilliporeSigma) or rabbit anti-PDGFRα (RRID: AB_3268408, Abcam) antibodies were coupled to magnetic beads (Dynabeads M-280 Tosylactivated, Invitrogen) according to the manufacturer’s protocol. Briefly, 17.5 µL of 30 mg/mL of dynabeads were washed several times in buffer A and were resuspended in buffer A containing 3 M ammonium sulfate. This solution was added to 12.5 µg of either CD24 or PSA-NCAM antibodies. They were incubated overnight at 37°C on a shaker, and the beads were precipitated for 1 min using a magnet. The antibody-coupled beads were washed with 1 mL of buffer D for 1 h at 37°C. The beads were precipitated with a magnet and underwent additional washes in buffer C. The antibody-coupled beads were resuspended in buffer C and were stored at 4°C. Immediately before the stereotactic injections, they were vortexed and sonicated.

#### Time-lapse imaging in acute brain slices

Acute brain slices were prepared as described previously^58^. Briefly, the mice were anesthetized with ketamine (100 mg/kg) and xylazine (10 mg/kg) and were perfused transcardially with modified oxygenated artificial cerebrospinal fluid (ACSF) containing (in mM): 210.3 sucrose, 3 KCl, 2 CaCl_2_.2H_2_O, 1.3 MgCl_2_.6H_2_O, 26 NaHCO_3_, 1.25 NaH_2_PO_4_.H_2_O, and 20 glucose. The brains were then quickly removed, and 250-µm-thick slices were cut using a vibratome (HM 650V; ThermoFisher Scientific). The slices were kept at 37°C in ACSF containing (in mM): 125 NaCl, 3 KCl, 2 CaCl_2_.2H_2_O, 1.3 MgCl_2_.6H_2_O, 26 NaHCO_3_, 1.25 NaH_2_PO_4_.H_2_O, and 20 glucose under oxygenation for no more than 6-8 h. The acute slices were transferred to the imaging chamber and were continuously superfused with ACSF at the rate of 1-1.5 mL/min. Ca^2+^ imaging was performed at 31-33_°_C using a BX61WI (Olympus) upright microscope equipped with a motorized *z* drive, a 40x water immersion objective (NA=0.8), a CCD camera (CoolSnap HQ), and mercury arc lamp. GCaMP6s was excited using a 482/35 nm excitation filter and emission was collected using a 536/40 nm filter. Imaging was performed every 30 s for 1 h with multiple z stacks (5-9 stacks at 3-µm intervals). In some experiments, time-lapse imaging was performed using an Axio Imager M2 upright microscope equipped with a motorized *z* drive, a Plan-Apo 40x water immersion objective (NA=1.0), an Axiocam 705 camera, and a Colibri 5 LED light source. For the pharmacological activation of Piezo1, we bath-applied Yoda 1 (5 μM, Tocris) and assessed Ca^2+^ activity in NSCs in acute adult brain slices derived from mice that had been electroporated with GFAP-GCaMP6s and Prom-mCherry at P1. We first recorded Ca^2+^ activity in NSCs under baseline conditions for 30 min, followed by a 45-min bath application of Yoda1. We compared the frequency and amplitude of Ca^2+^ events during 30 min of baseline recording with the those of the last 30 min of Yoda1 application.

For time-lapse imaging of CD24 antibody-coupled beads, a binning of 4x4 was used to increase the temporal resolution and the “stream acquisition” feature of Methamorph software in a single z-section. This roughly results in an acquisition rate of 80-100 frames per second. The imaging was performed for 1-2 min. The same parameters were used for time-lapse imaging Lck-GFP-electroporated ECs with a 5-10 ms excitation time. In some experiments, an upright spinning disk confocal microscope (Nikon) equipped with a 25x water immersion objective (Nikon, NA=1.1) and a Yokogawa CSU-X1 disk was used. GFP and magnetic beads, which are fluorescent at 561 nm, were excited with 488 and 561 solid state lasers and the emission was captured using a Prime BSI-Express camera after passing through the appropriate emission and dichroic filters. This imaging system made it possible to further increase the temporal resolution of the imaging up to 180 frames per second using a 5-ms excitation time and 2x2 binning.

#### Ca^2+^ imaging and EC cilia beating analyses

Ca^2+^ responses were analyzed using a custom-written script in MATLAB (MathWorks Inc., USA), as described previously^34^. Briefly, the regions of interest (ROI) were manually drawn around the soma of the cell to extract the GCaMP6s fluorescence signal. Ca^2+^ activity was calculated as relative changes in the percentage of ΔF/F = (F-F_back_)/F_back_, where F is the GCaMP6 intensity in the ROI and F_back_ is the background signal. A multiple threshold algorithm was used to analyze spontaneous Ca^2+^ activity in NSCs^59^, as described previously^34^. Briefly, the mean standard deviation (SD) of each Ca^2+^ trace was first calculated, and all peaks with an amplitude greater than 2 times the SD were measured. These peaks were then removed from the Ca^2+^ traces, and the mean SD of the Ca^2+^ trace was re-calculated to depict events greater than 2 times the new SD. This procedure was repeated one more time, which allowed us to depict all Ca^2+^ events regardless of their amplitudes. Three different thresholds for the detection of Ca^2+^ events are shown in the **Figures** by dashed lines.

A custom-written script in MATLAB (MathWorks Inc., USA) was used to analyze the cilia beating of ECs labeled either fluorescently with Lck-GFP or coupled with beads. As for the Ca^2+^ analysis, the images were recorded for the XY positions and were corrected for xy drift using the StackReg ImageJ plugin, as required. Our analysis was focused on tracking single beads or fluorescently labeled single cilia of multiciliated ECs that can be reliably tracked in multiple time-points of 1-2-min. For the bead analyses, the regions containing single beads were selected and the background was subtracted based on the difference in the intensities of the pixels of the signal (beads) and the background. The beads were detected based on multiple high-intensity pixels, and the *xy* position of the centroid of the signal (beads) was calculated in each frame of a time-lapse movie (**Figure 1G**). This made it possible to plot bead displacement over time (**Figure 1G**) and to calculate the frequency. The same script was used for the cilia analyses, with slight modifications. Instead of the centroid of the bead, the displacement of the tip of individual EC cilia was tracked.

#### *Post hoc* identification of imaged cells

For the *post hoc* identification of cells imaged in acute brain slices, 250-µm-thick slices were fixed in 4% PFA overnight and were pre-permeabilized with methanol and acetone (30 min each at – 30°C) prior to immunostaining. The brain slices were incubated with anti-GFP to reveal GFAP-GCaMP6s labeling as well as with anti-Ki67 and anti-mCherry primary antibodies and were processed using secondary antibody labeling and confocal imaging, as described above. Imaged cells that were not retrieved by the *post hoc* identification analysis were discarded.

#### Electro-magnetic stimulation in acute brain slices

Electro-magnetic stimulation was used to modulate EC cilia beating *ex vivo* in acute brain slices. Acute brain slices were prepared from mice injected with CD24 antibody-coupled beads, as described above. The electrode, which consisted of a coiled copper wire looped at the tip (**Figure 1I**), was lowered and positioned approximately 50-75 µm from the surface of the slice. The wire was approximately 150 µm in diameter and the looped part was bent parallel to the surface of the slice. A 5-s pulse of a 9.5-A current was applied from a current source (power supply 1-60 V, 15-A switch, B&K Precision) resulting in a 2.8mT magnetic field measured with IDR-309 gaussmeter (Integrity Design & Research)at the tip of the electrode. Different current magnitudes, pulse durations, modes of application (ramp or step-wise pulse), and copper wire diameters were empirically tested to determine the efficiency of the magnetic field for stopping the beads.

#### Magnetic stimulation *in vivo*

To modulate CD24 antibody-coupled magnetic beads *in vivo*, a custom-made behavioral chamber consisting of two 33x19x14-cm cages connected by a 24.2-cm hamster tunnel purchased from a local pet-shop was used. We installed permanent 4x1x1/4-in 1.4T neodymium magnets (grade N52, CMS Magnetics) on both sides of the tunnel, which resulted in a 300-350-mT magnetic field inside the tunnel, which was measured with gaussmeter (Integrity Design & Research). No magnetic fields were detected inside the rat cages. It should be noted that in contrast to *ex vivo* preparation where the electro-magnetic field was given very close to the magnetic beads, in our *in vivo* experiments, we used permanent magnets that required greater strength to affect bead movement several centimeters away. This explains the differences in the strengths of magnetic field applied *ex vivo* and *in vivo*. The mice were injected in the LV with CD24 antibody-coupled magnetic beads under isoflurane anesthesia at the following coordinates (with respect to the bregma): AP 0.3 mm, ML 0.9 mm, and DV 2.7 and 2.2 mm. Immediately after recovery from the stereotactic (usually 20-30 min), the mice were placed in the behavioral chamber and were allowed to explore it freely for 3 h. Behavioral cameras were used to record the animals’ behavior and to calculate the number of tunnel crosses and the overall time that animals spent in the tunnel. To assess how the exposure of the mice to behavioral chambers equipped with permanent magnets affected EC cilia beating, acute brain slices from mice injected with CD24 antibody-coupled beads were prepared immediately after the 3-h period and bead-coupled cilia movement was imaged, as described above. As a control, slices from mice that had been injected with magnetic beads coupled to CD24 antibodies were exposed to behavioral chambers without the permanent magnets. Time-lapse imaging was performed for up to 1 h after the preparation of the brain slices to minimize any possible washout effect of magnetic stimulation.

Following the establishment of the protocol for modulating EC cilia beating *in vivo* and validating that it effectively inhibits cilia beating, how the inhibition of EC cilia beating affects NSC quiescence/activation dynamics was then assessed. To do so, mice previously electroporated with GFAP-GFP and Prom-mCherry plasmids at P0-P1 were intraventricularly injected with CD24 antibody-coupled beads at P45-P60 and were allowed to explore the behavioral chambers connected by a tunnel with permanent magnets. As controls, we used two groups of mice that had been electroporated with the GFAP-GFP and Prom-mCherry plasmids at P0-P1 and were either injected with magnetic beads coupled to PSA-NCAM antibodies (PSA-NCAM is not present on the cilia of ECs and thus do not result in beads binding to the cilia (see **Figure 1B**)) or mice injected with magnetic beads coupled with CD24 antibodies but exposed to behavioral chambers without permanent magnets. All three groups of mice were allowed to freely explore the behavioral chambers for 3 h. The total time spent in the tunnel and the number of tunnel crossings were video-recorded and analyzed offline. Immediately after the 3-h period, the mice were perfused transcardially with 0.9% NaCl followed by 4% PFA, and 40-µm-thick sagittal sections were processed for GFP, mCherry, and Ki67 immunolabeling. The percentage of activated NSCs (GFAP-GFP+/Prom-mCherry+/Ki67+) was counted along the entire lateral wall of the LV. Quantification was performed manually using the ImageJ cell counter plugin.

To assess whether the inhibition of EC cilia beating affects global cerebrospinal fluid (CSF) flow, we performed *in vivo* imaging of Alexa488-dextran (10kDa) dye diffusion following its injection into one hemisphere. CD1 mice were injected with CD24 antibody coupled beads into the LV as described above and were either exposed or not exposed to the permanent magnet for 3 h in the behavioral chambers. Immediately after 3h-period, the mice were injected with 1 μl of Alexa488-dextran into the LV of the same hemisphere and the spread of the dye was assessed in both hemispheres during 90 min using Newton FT-500 *in vivo* imaging system, using following parameters: Ultimate 4, aperture 1.4, and exposure time 100 ms.

#### Live tissue passive microrheology measurements and processing

The cytoplasmic dynamics of NSCs expressing or lacking TRPM3 were quantified using an organelle-tracking–based microrheology approach as previously described^38^. Briefly, the acute adult brain slices derived from mice that had been previously electroporated either with *LacZ* or *TrpM3* gRNAs were prepared as described above and were incubated for 1 h with 100 nM of LysoTracker™ DeepRed. The fluorescently labeled lysosomes (LysoTracker™) were imaged using an upright spinning disk confocal microscope (Nikon) equipped with a 25x water immersion objective (Nikon, NA=1.1) and a Yokogawa CSU-X1 disk. GFP and mCherry were excited with 488 and 561 solid state lasers, while LysoTracker™ DeepRed was excited at 640 nm. The emission was captured using a Prime BSI-Express camera after passing through the appropriate emission and dichroic filters. After ensuring that labeled lysosomes were present in NSCs electroporated with GFAP-GFP and either *Trpm3* gRNAs (pU6-*Trpm3* gRNA-Prom-Cas9-T2A-mCherry) or *LacZ* (control) gRNAs, we imaged a single optical section using 2x2 binning and an acquisition rate of 75 frames per second. This high temporal resolution made it possible to capture short-timescale intracellular fluctuations.

For lysosome tracking, the time-lapse imaging stacks were first corrected for sample drift using the Fast4DReg algorithm implemented in Fiji/ImageJ^60,61^, which performs sub-pixel 3D registration by phase correlation in Fourier space. The resulting transformation matrices were applied to all channels prior to tracking. Lysosomes trajectories were extracted using the TrackMate plugin in Fiji^62^ with sub-pixel localization accuracy. Spots were detected using a Difference-of-Gaussian detector (estimated diameter = 1 µm), and trajectories were linked across frames using the linear assignment problem (LAP) tracker. Only trajectories longer than 500 frames were retained for analysis. and were linked across frames using a linear assignment (LAP) tracker. Trajectory data were exported as XML files and were analyzed in MATLAB (MathWorks Inc., USA) using custom scripts.

For each trajectory, the root mean squared displacement (RMSD) as a function of lag time *𝜏𝜏* was computed in MATLAB as

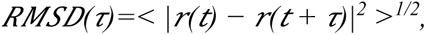

where <> denotes averaging over time t. A representative *RMSD(𝜏)* curve is shown in **Supp. Figure 2**. As expected, *RMSD(𝜏)* exhibited a short-time plateau, which is consistent with thermal equilibrium at high frequencies. We defined RMSD_0_ = *RMSD(𝜏=12ms)*, corresponding to the highest imaging frequency as a measure of the plateau value, which is proportional to cytoplasmic mechanical compliance at thermal equilibrium.

#### RNAscope *in situ* hybridization

RNAscope Multiplex Fluorescent Reagent kits (Advanced Cell Diagnostics) were used according to the manufacturer’s protocol for the *in situ* hybridization of Trpm3, GFP, and mCherry. *In situ* hybridization was performed on 15-µm-thick fresh frozen brain slices from CD1 mice electroporated with GFAP-GFP and Prom-mCherry at P0-P1, as described previously^34,35^. Briefly, the frozen brain slices were mounted on slides and were dried for 1 h at –20 °C. They were then incubated in cold 4% PFA for 15 min and were sequentially dehydrated in 50%, 70%, and 100% ethanol solutions for 5 min each at RT. The slides were treated with Protease IV for 30 min at RT for antigen retrieval. The probes were diluted (1:50) and were applied to the slides for 2 h at 40°C. The slides were then incubated with a series of preamplifier and amplifier reagents at 40°C (AMP1 for 30 min, AMP2 for 15 min, AMP3 for 30 min, and AMP4 Alt A for 15 min) according to the manufacturer’s protocol. Images were acquired using an inverted Zeiss microscope (LSM 700, AxioObserver) with a 20x air immersion objective (NA: 0.9) or a 63x oil immersion objective (NA: 1.4).

#### Nuclei isolation, snRNA-Seq library construction and sequencing

Nuclei were isolated from 2mmx2mmx5mm pieces of collected and snap frozen tissue according to the manufacturer’s recommended protocol (10X Genomics nuclear isolation kit). Single nuclei from each sample (4,000 for individual pilot or 25,000 for pooled samples) were loaded for partitioning using 10X Genomics NextGEM Gel Bead emulsions (v3.1). Each sample was processed according to the manufacturer’s recommended protocol (PCR amplification steps were run at 14X, and 14X respectively). Final cDNA library size determination and QC was performed using Agilent TapeStation D1000 assay. Sequencing was performed using Illumina NovaSeq S2 and SP 100 cycle dual lane flow cells over multiple rounds at the University of Calgary Centre for Health Genomics and Informatics (CHGI). A total of 44,676 nuclei were captured across all samples. Each sample was sequenced to approximately 20,000 reads per nuclei. Sequencing reads were aligned using CellRanger v7.1.0 pipeline to the standard pre-built mouse reference genome (mm10). Samples that passed alignment QC were aggregated using CellRanger aggr with between-sample normalization to ensure each sample received an equal number of mapped reads per cell in the final dataset.

Since *cellranger* indicated a high amount of ambient RNA, ambient RNA was removed using *CellBender*^63^. The six samples were then combined in *scanpy*^64^. Nuclei were kept if they had a minimum of 200 genes and between 1000 and 150,000 counts. Genes were kept if they were present in at least 5 nuclei. An overview of raw quality control (QC) statistics can be found in **Table 1**. Moreover, nuclei with more than 20 percent mitochondrial reads were removed. Potential doublets were removed using *scrublet* with a threshold of 0.2^65^. This yielded a count matrix of 31,307 nuclei and 26,147 genes. Library size normalization was done using *scran* as described^66,67^. Leiden clustering at a resolution of 0.5 resulted in 13 clusters, of which clusters 5, 6, 10 and 11 were further subclustered for a better cell type separation of TAP, Neuroblasts, endothelial cells, pericytes, and the oligodendrocyte lineage. The resulting 19 clusters were annotated into 12 cell types using marker genes from literature. Additional sub-clustering of Astrocytes/NSCs, Ependymal cells, Neuroblasts and TAPs allowed to separate aNSCs from qNSCs. Differential expression analysis was done using *sc.tl.rank_genes_groups*. As differential expression analysis showed some mitochondrial transcripts, that were most likely contaminants in the library prep, we opted to remove them before performing Gene Ontology (GO) analysis for Biological Processes. GO analysis was done in *ShinyGo v0.85* with default parameters against the mouse genome^68^.

**Table 1:** Overview of raw QC statistics.

| Sample | Condition | Mean Counts | Mean percentage mitochondrial reads |
| --- | --- | --- | --- |
| 1 | CD24 beads + magnet | 6074 | 5.55 |
| 2 | CD24 beads, no magnet | 6823 | 2.75 |
| 3 | CD24 beads, no magnet | 3404 | 4.23 |
| 4 | CD24 beads + magnet | 3625 | 7.09 |
| 5 | CD24 beads, no magnet | 3149 | 7.55 |
| 6 | CD24 beads + magnet | 4051 | 2.79 |

#### Osmotic mini-pump installation

CD1 mice previously electroporated with GFAP-GFP and Prom-mCherry plasmids at P0-P1 were anesthetized at P45-P60 with isoflurane to inject CD24 antibody-coupled magnetic beads and to install osmotic minipumps during the same surgery. All-plastic PEEK-based cannulas (HRS Scientific) connected to the osmotic minipumps (model 2001D, infusion rate 8 uL/h Alzet) were used. Immediately after the intraventricular injection of the CD24 antibody-coupled magnetic beads (as described above), the cannula was stereotactically implanted at the following coordinates (with respect to the bregma): AP 0.3 mm, ML 0.9 mm, and DV ∼1.9mm (length of the cannula). The mice were maintained at 37_°_C throughout the surgery. The cannula was connected to an osmotic minipump filled either with vehicle (0.9% NaCl; Sigma) or 2.5 µM CIM0216 (Tocris), a selective TRPM3 agonist. Dental cement (Metabond, Parkell) was used to secure the cannulae to the skull, and the skin was sutured. Immediately after recovery from the stereotaxic surgery (30-45 min), the mice were placed in the behavioral chambers connected with a tunnel with permanent neodymium magnets on both sides of the tunnel. They were allowed to freely explore the behavioral chambers for 3 h, and the time in the tunnel and the number of tunnel crosses were analyzed offline. The mice were perfused transcardially with 0.9% NaCl followed by 4% PFA and were processed for GFP, mCherry, and Ki67 immunolabeling. The percentage of activated Ki67+ NSCs (GFAP-GFP+/Prom-mCherry+) was determined as described.

## QUANTIFICATION AND STATISTICAL ANALYSIS

Data are expressed as means ± SEM. The individual values of all experiments are also indicated in the text and corresponding figure legends. Ca^2+^ activity was determined prior to the *post hoc* identification of imaged cells. As such, the investigator was unaware of whether it reflected the activity of qNSCs or aNSCs. Statistical significance was determined using a paired or an unpaired two-sided Student’s *t*-test or one-way ANOVA followed by an LSD-Fisher *post hoc* test using Statistica software. Equality of variance for the unpaired t-test was verified using an F-test. The exact values of *n* and its representation (cells, animals) for all experiments are indicated in the corresponding figure legends. The levels of significance were \**p*<0.05, \*\**p*<0.01, \*\*\**p*<0.001 and \*\*\*\**p*<0.0001.

