## Supplementary Figures for "Cilia beating of ependymal cells regulates adult neural stem cell quiescence via mechanical forces mediated by PKD1/2-TRPM3"

### Supplemental Information

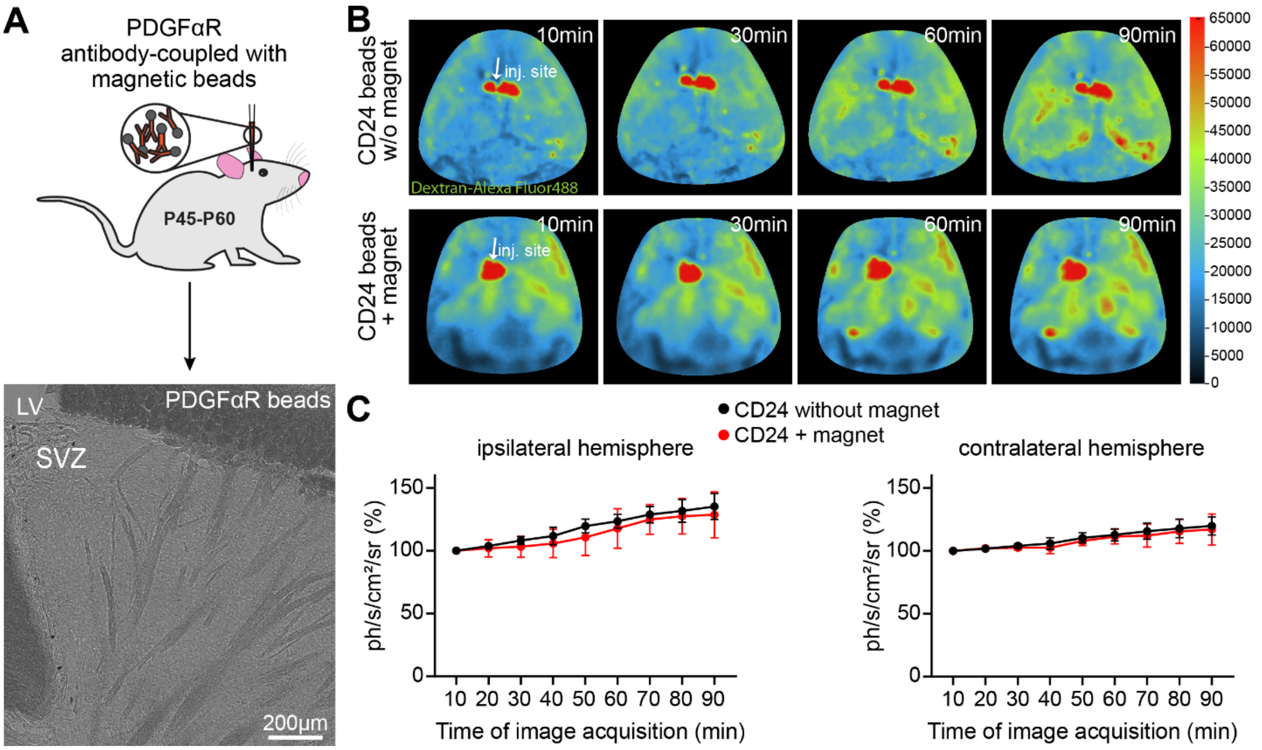


**Supplementary Figure 1. Inhibition of EC cilia beating does not affect global CSF flow.**

**A:** Experimental approach and sagittal section of adult mouse brains showing the lack of binding of beads coupled to PDGFRα antibodies to ECs lining the wall of the LV. **B:** *In vivo* imaging of Alexa488-dextran (10 kDa) diffusion following inhibition of EC cilia beating. The mice were injected with CD24 antibody coupled magnetic beads and were either exposed or not exposed to the permanent magnet by placing them into the behavioral chambers for 3 h. Immediately after 3h-period fluorescent dextran was injected into the hemisphere injected with CD24 antibody coupled beads and its spread was assessed for 90 min. **C:** The quantification of the spread of fluorescent dextran in the injected, ipsilateral (left) and contralateral (right) hemispheres. Note that inhibition of EC cilia beating does not affect the spread of fluorescent dextran, indicating for a normal CSF flow. N = 3 mice per group.

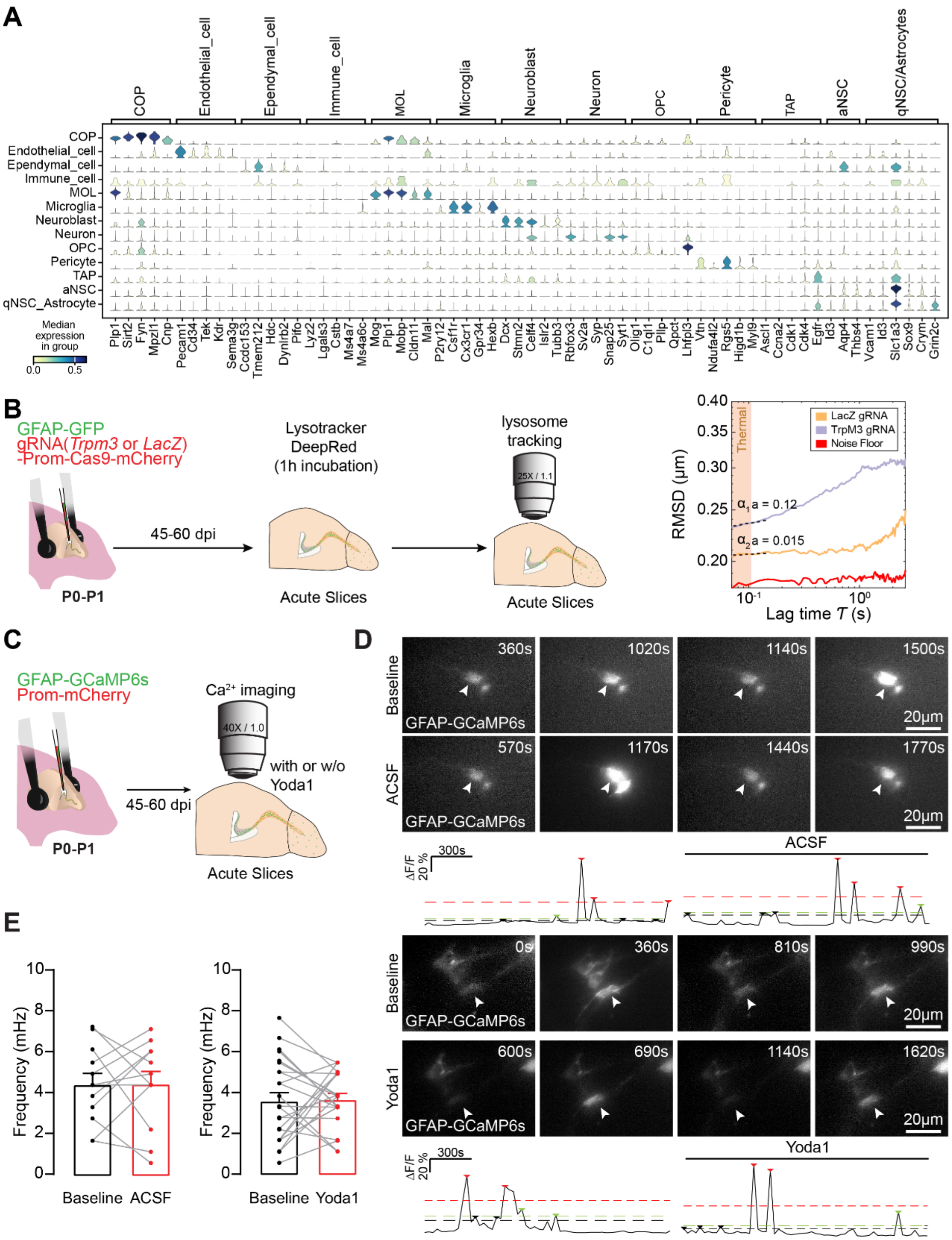


**Supplementary Figure 2. snRNA-seq analysis and the role of mechano-sensitive receptors.**

**A: The** cell types of snRNA-seq analysis defined based on marker gene expression. **B:** Schematic representation of the experimental approach to assess the role of TRPM3 in the intracellular stiffness. The right panel shows the root mean-squared displacement (RMSD, y axis, log scale) of fluorescent lysosomes tracked in the soma of LacZ control cells (orange) and TRPM3 knockdown cells (purple), plotted against lag time $\tau$ (x axis, log scale). The shaded area below 0.1 marks the short-time regime dominated by local viscoelastic mechanics. Dashed lines indicate log–log fits giving anomalous diffusion exponents $\alpha = 0.015$ for LacZ and $\alpha = 0.12$ for TrpM3 KD. **C:** Schematic diagram of the experimental approach to assess the role of Yoda1, a Piezo1 activator, on $Ca^{2+}$ activity in NSCs. **D:** Representative images and traces of $Ca^{2+}$ activity in NSCs under control condition and following bath-application of Yoda1. **I**: Quantification of the frequency and amplitude of $Ca^{2+}$ events in NSCs under control conditions and following Yoda1 application.
